# PTPN2 deletion in T cells promotes anti-tumour immunity and CAR T cell efficacy in solid tumours

**DOI:** 10.1101/757419

**Authors:** Florian Wiede, Kun-Hui Lu, Xin Du, Shuwei Liang, Katharina Hochheiser, Garron T. Dodd, Pei Goh, Conor Kearney, Deborah Meyran, Paul A. Beavis, Melissa A. Henderson, Simone L. Park, Jason Waithman, Sheng Zhang, Zhong-Yin Zhang, Jane Oliaro, Thomas Gebhardt, Phillip K. Darcy, Tony Tiganis

## Abstract

Although adoptive T cell therapy has shown remarkable clinical efficacy in hematological malignancies, its success in combating solid tumours has been limited. Here we report that PTPN2 deletion in T cells enhances cancer immunosurveillance and the efficacy of adoptively transferred tumour-specific T cells. T cell-specific PTPN2 deficiency prevented tumours forming in aged mice heterozygous for the tumour suppressor p53. Adoptive transfer of PTPN2-deficient CD8+ T cells markedly repressed tumour formation in mice bearing mammary tumours. Moreover, PTPN2 deletion in T cells expressing a chimeric antigen receptor (CAR) specific for the oncoprotein HER-2 increased the activation of the Src family kinase LCK and cytokine-induced STAT-5 signalling thereby enhancing both CAR T cell activation and homing to CXCL9/10 expressing tumours to eradicate HER-2+ mammary tumours *in vivo*. Our findings define PTPN2 as a target for bolstering T-cell mediated anti-tumour immunity and CAR T cell therapy against solid tumours.

---

Tumours can avoid the immune system by co-opting immune checkpoints to directly or indirectly inhibit the activation and function of cytotoxic CD8+ T cells ^1, 2^. In particular, the inflammatory tumour microenvironment can upregulate ligands for T cell inhibitory receptors such as programmed cell death protein-1 (PD-1) on tumour cells to inhibit T cell signalling and promote the tolerisation or exhaustion of T cells ^1, 2^. Immune checkpoint receptors, including PD-1 and cytotoxic T-lymphocyte antigen-4 (CTLA-4) can suppress the amplitude and/or duration of T cell responses by recruiting phosphatases to counteract the kinase signalling induced by the T cell receptor (TCR) and co-stimulatory receptors such as CD28 on αβ T cells ^2, 3^.

Protein tyrosine phosphatase N2 (PTPN2) negatively regulates αβ TCR signalling by dephosphorylating and inactivating the most proximal tyrosine kinase in the TCR signalling cascade, the Src family kinase (SFK) LCK ^4, 5^. PTPN2 also antagonises cytokine signalling required for T cell function, homeostasis and differentiation by dephosphorylating and inactivating Janus-activated kinase (JAK)-1 and JAK-3 and their target substrates signal transducer and activator of transcription (STAT)-1, STAT-3 and STAT-5 in a cell context-dependent manner ^6–9^. By dephosphorylating LCK PTPN2 sets the threshold for productive TCR signalling and prevents overt responses to self-antigen in the context of T cell homeostasis and antigen cross-presentation to establish peripheral T cell tolerance ^10, 11^. The importance of PTPN2 in T cells in immune tolerance is highlighted by the development of autoimmunity in aged T cell-specific PTPN2-deficient mice on an otherwise non-autoimmune C57BL/6 background ^5^, the systemic inflammation and autoimmunity evident when PTPN2 is deleted in the hematopoietic compartment of adult C57BL/6 mice ^8^ and the accelerated onset of type 1 diabetes in T cell-specific PTPN2-deficient mice on the autoimmune-prone non-obese diabetic (NOD) background ^12^. In humans, *PTPN2* deficiency is accompanied by the development of type 1 diabetes, rheumatoid arthritis and Crohn’s disease ^13, 14^. The autoimmune phenotype of PTPN2-deficient mice is reminiscent of that evident in mice in which the immune checkpoint receptors PD-1 ^15–17^ or CTLA-4 ^18, 19^ have been deleted. Whole-body PD-1 deletion results in spontaneous lupus-like autoimmunity in C57BL/6 mice ^15^ and accelerated type 1 diabetes onset in NOD mice ^17^, whereas CTLA4 deletion in C57BL/6 mice results in marked lymphoproliferation, autoreactivity and early lethality ^18, 19^. Although PD-1 and/or CTLA4 blockade can be accompanied by the development of immune-related toxicities, antibodies targeting these receptors have nonetheless shown marked therapeutic efficacy in various tumours, including melanomas, non-small cell lung carcinomas, renal cancers and Hodgkin lymphoma ^1, 2^. Accordingly, we sought to assess the role of PTPN2 in T cell-mediated immunosurveillance and the impact of targeting PTPN2 on adoptive T cell immunotherapy. We especially focused on CAR T cell therapy, which has shown marked clinical efficacy in B cell acute lymphoblastic leukemia (ALL), but has been largely ineffective in solid tumours ^20–23^.

## RESULTS

### PTPN2 deletion prevents tumour formation in p53^+/–^ mice

First we determined the impact of deleting PTPN2 in T cells on tumour formation in mice heterozygous for *p53*, the most commonly mutated tumour suppressor in the human genome ^24^. In humans, inheritance of one mutant allele of p53 results in a broad-based cancer predisposition syndrome known as Li-Fraumeni syndrome ^25^. In mice, *p53* heterozygosity results in lymphomas and sarcomas, as well as lung adenocarcinomas and hepatomas in 44% of mice by 17 months of age with the majority of tumours exhibiting *p53* loss of heterozygosity (LOH) ^26^. We crossed control (*Ptpn2^fl/fl^*) and T cell-specific PTPN2-null mice (*Lck*-Cre;*Ptpn2^fl/fl^*) onto the *p53^+/–^* background and aged the mice for one year. Upon necropsy 15/28 (54%) *Ptpn2^fl/fl^*;*p53^+/–^* mice developed various tumours including thymomas, lymphomas, sarcomas, carcinomas and hepatomas (Fig. 1a; Fig. S1; Table S1) as reported previously for *p53* heterozygous mice ^26^. In addition, 6/28 mice exhibited splenomegaly accompanied by the accumulation of CD19^+^IgM^hi^CD5^hi^B220^int^ B1 cells consistent with the development of B cell leukemias (Fig. 1b), whereas CD3 negative CD4+CD8+ double positive cells reminiscent of T cell leukemic blasts (FSC-A^hi^) were evident in the thymi or peripheral lymphoid organs of 5/28 mice (Fig. 1a-b; Table S1). Histological analysis revealed disorganised thymic, lymph node or splenic tissue architecture in diseased *Ptpn2^fl/fl^*;*p53^+/–^* mice that was predominated by larger lymphoblasts consistent with the accumulation of pre-leukemic/leukemic cells (Fig. S1). By contrast no *Lck*-Cre;*Ptpn2^fl/fl^*;*p53^+/–^* mice (0/22) developed any overt tumours, splenomegaly or abnormal lymphocytic populations as assessed by gross morphology or flow cytometry and lymphoid organ tissue architecture was normal (Fig. 1; Fig. S1). PTPN2 deficiency in T cells can result in inflammation/autoimmunity in aged C57BL/6 mice ^5^. Accordingly, we determined if PTPN2-deficiency might exacerbate inflammation in *p53^+/–^* mice. We found that inflammation, as assessed by measuring the proinflammatory cytokines IL-6, TNF and IFNγ in plasma, was elevated in *Lck*-Cre;*Ptpn2^fl/fl^*;*p53^+/–^* mice (Fig. S2a), as seen in aged *Lck*-Cre;*Ptpn2^fl/fl^* mice (Fig. S2b), but this did not exceed that occurring in *Ptpn2^fl/fl^*;*p53^+/–^* littermate controls. Aged *Lck*-Cre;*Ptpn2^fl/fl^*;*p53^+/–^* mice also had lymphocytic infiltrates in their livers (Fig. S2c), forming what resembled ectopic lymphoid-like structures ^27^ and this was accompanied by liver damage and ensuing fibrosis (Fig. S2c). However lymphocytic infiltrates and fibrosis were also evident in the livers of tumour-bearing *Ptpn2^fl/fl^*;*p53^+/–^* mice (Fig. S2c). Taken together results indicate that PTPN2 deficiency in T cells can prevent the formation of tumours induced by *p53* LOH without exacerbating inflammation.

**Fig. 1.**
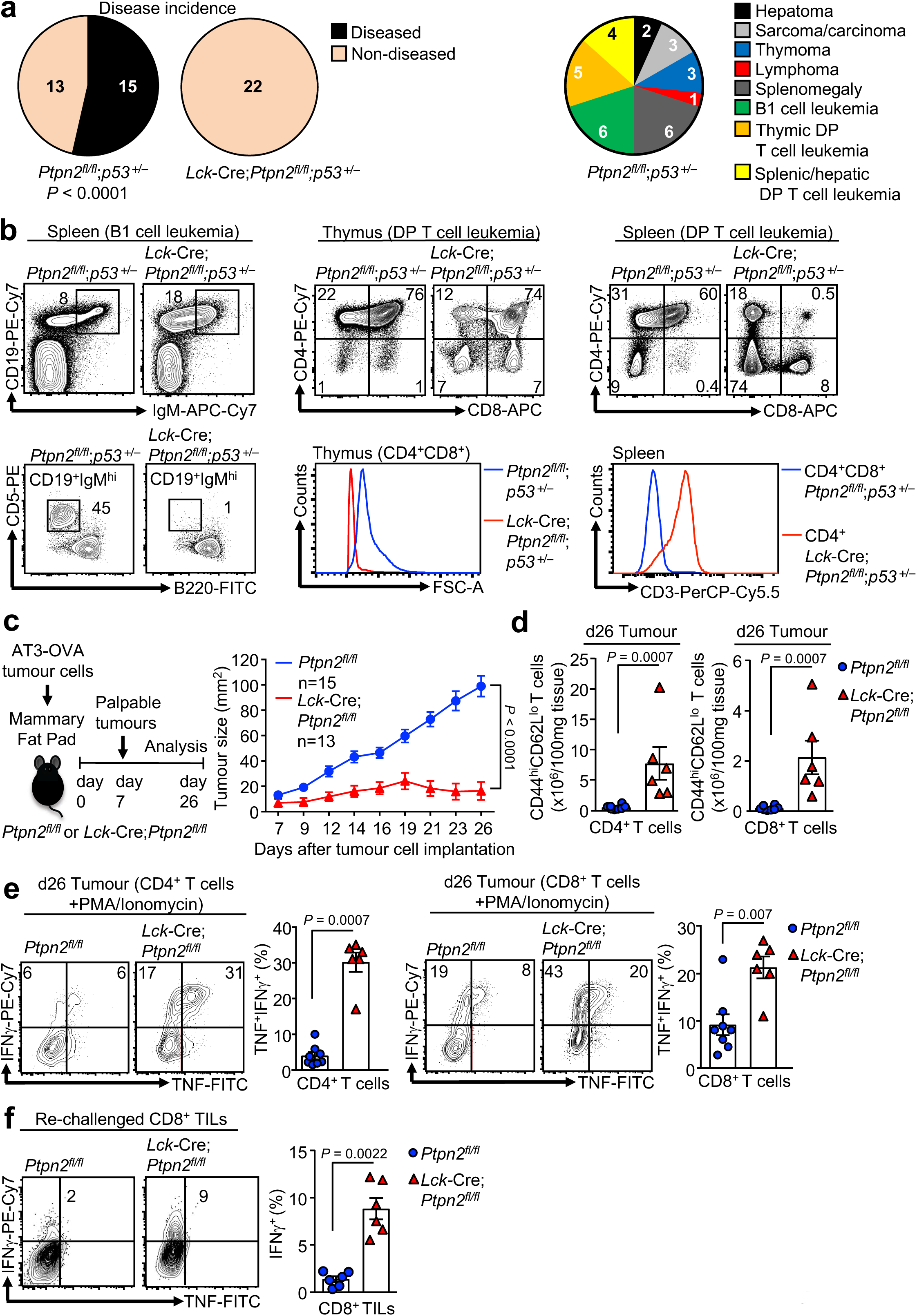
PTPN2-deletion in T cells increases tumour immunosurevillance. **a-b)** 12 month old *Ptpn2^fl/fl^*;*p53^+/–^* and *Lck*-Cre;*Ptpn2^fl/f^;p53^+/–^* mice were assessed for **a**) disease and tumour incidence. **b**) Lymphocytes from 12 month old *Ptpn2^fl/fl^*;*p53^+/–^* and *Lck*-Cre;*Ptpn2^fl/f^;p53^+/–^* mice were stained for CD19, B220, IgM, CD5, CD3, CD4 and CD8 and analyzed by flow-cytometry. Representative flow cytometry profiles are shown. Significance in (a) was determined using two-sided Fisher’s exact test. **c**) AT-3-OVA breast cancer cells (5×10^5^) were injected into the fourth inguinal mammary fat pads of female *Ptpn2^fl/fl^* and *Lck*-Cre;*Ptpn2^fl/fl^* mice and tumour growth monitored over 26 days. **d**) At day 26 (d26) tumour infiltrating lymphocytes (TILs) were stained for CD4, CD8, CD62L and CD44 and T cell numbers were determined. **e**) d26 TILs were stained for CD4, CD8 and intracellular IFNγ and TNF and the proportion of IFNγ^+^ versus IFNγ^+^TNF^+^ T cells determined. **f**) d26 TILS were incubated with AT-3-OVA tumour cells isolated from tumour bearing C57BL/6 mice and stained for CD8 and intracellular IFNγ and TNF and the proportion of IFNγ^+^ T cells determined. Representative flow cytometry profiles and results (means ± SEM) from at least two independent experiments are shown. In (c) significance was determined using 2-way ANOVA Test and in (d-f) significance determined using 2-tailed Mann-Whitney U Test.

### PTPN2 deficiency enhances T cell-mediated immunosurveillance

At least one mechanism by which PTPN2-deficiency might prevent tumour formation in *p53^+/–^* mice might be through the promotion of T cell-mediated tumour immunosurveillance. To explore this we first assessed the growth of syngeneic tumours arising from ovalbumin (OVA)-expressing AT-3 (AT-3-OVA) adenocarcinoma cells implanted into the inguinal mammary fat pads of *Ptpn2^fl/fl^* versus *Lck*-Cre;*Ptpn2^fl/fl^* C57BL/6 mice (Fig. 1c); AT-3 cells lack estrogen receptor, progesterone receptor and ErbB2 expression and are a model of triple negative breast cancer ^28^^29^. Whereas AT3-OVA cells grew readily in *Ptpn2^fl/fl^* mice, tumour growth was markedly repressed in *Lck*-Cre;*Ptpn2^fl/fl^* mice so that tumour progression was prevented in 5/13 mice and eradicated in 2/8 of the remaining mice after tumours had developed. The repression of tumour growth was accompanied by the infiltration of CD4+ and CD8+ effector/memory (CD44^hi^CD62L^lo^) T cells into tumours (Fig. 1d). Consistent with our previous studies ^5^, PTPN2-deficient CD25^hi^FoxP3^+^ regulatory T cells (T_regs_) were increased rather than decreased in AT3-OVA tumours (Fig. S2d) and their activation was increased (data not shown) precluding the repression of tumour growth being due to defective Treg-mediated immunosuppression. Moreover, tumour-infiltrating PTPN2-deficient CD4+ and CD8+ effector/memory T cells were significantly more active, as assessed by the PMA-Ionomycin-induced production of markers of T cell cytotoxicity *ex vivo*, including interferon (IFN) γ and tumour necrosis factor (TNF) (Fig. 1e); by contrast splenic PTPN2-deficient T cells exhibited comparatively small increases in cytotoxic capacity as assessed by IFNγ/TNF expression (data not shown). To directly assess the influence of PTPN2-deficiency on T cell-mediated immunosurveillance, we next isolated tumour-infiltrating CD8+ T cells from *Ptpn2^fl/fl^* versus *Lck*-Cre;*Ptpn2^fl/fl^* mice and assessed their activation by measuring IFNγ production *ex vivo* upon re-challenge with tumour cells isolated from AT3-OVA tumours that had developed in *Ptpn2^fl/fl^* mice (Fig. 1f). *Ptpn2^fl/fl^* tumour-infiltrating CD8+ T cells remained largely unresponsive when re-challenged, consistent tolerisation. By contrast, PTPN2-deficient T cells exhibited significant increases in IFNγ and TNF consistent with increased effector activity (Fig. 1f). These findings point towards PTPN2 having an integral role in T cell-mediated immune surveillance.

To explore the cellular mechanisms by which PTPN2-deficiency might influence immunosurveillance, we determined whether PTPN2 deletion might promote the tumour-specific activity of adoptively transferred TCR transgenic CD8+ T cells expressing the OT-1 TCR specific for the ovalbumin (OVA) peptide SIINFEKL. Naive OT-1 T cells can undergo clonal expansion and develop effector function when they engage OVA-expressing tumours, but thereon leave the tumour microenvironment, become tolerised and fail to control tumour growth ^30–32^. The eradication of solid tumours by naive CD8+ T cells is dependent on help from tumour-specific CD4+ T cells ^31, 33^. Our previous studies have shown that PTPN2-deficiency enhances TCR-instigated responses and negates the need for CD4+ T cell help in the context of antigen cross-presentation ^11^. Accordingly, we determined whether PTPN2 deficiency might overcome tolerisation and render naive OT-1 CD8+ T cells capable of suppressing the growth of OVA-expressing tumours. To this end, naive OT-1;*Ptpn2^fl/fl^* or OT-1;*Lck*-Cre;*Ptpn2^fl/fl^* CD8+ T cells were adoptively transferred into immunocompetent and non-irradiated congenic C57BL/6 hosts bearing syngeneic tumours arising from AT-3-OVA cells inoculated into the mammary fat pad (**Fig. 2a**). As expected ^30, 31^ adoptively transferred naive (CD44^lo^CD62L^hi^) *Ptpn2^fl/fl^* OT-1 CD8+ T cells (Gating strategy; Fig. S3) had no overt effect on the growth of AT-3-OVA mammary tumours when compared to vehicle-treated tumour-bearing mice (**Fig. 2a**). By contrast 5 days after adoptive transfer *Lck*-Cre;*Ptpn2^fl/fl^* OT-1 T cells completely repressed tumour growth (**Fig. 2a**). The repression of tumour growth was accompanied by an increase in *Lck*-Cre;*Ptpn2^fl/fl^* OT-1 T cells in the draining lymph nodes of the tumour-bearing mammary glands (Fig. S4a) and a marked increase in tumour-infiltrating *Lck*-Cre;*Ptpn2^fl/fl^* OT-1 T cells (**Fig. 2b**; Fig. S4b). At 9 days post adoptive transfer both tumour and draining lymph node *Lck*-Cre;*Ptpn2^fl/fl^* OT-1 T cells were more active, as assessed by the PMA/ionomycin-induced expression of effector molecules, including IFNγ, TNF and granzyme B (**Fig. 2c**; Fig. S4c). Although the expression of the T cell inhibitory receptors PD-1 and Lag-3 on tumour-infiltrating PTPN2-deficient OT-1 T cells at 9 days post-transfer was not altered (Fig. S4d), by 21 days post-transfer relative PD-1 and LAG-3 levels were reduced and CD44 was increased on PTPN2-deficient tumour-infiltrating and draining lymph node OT-1 T cells when compared to *Ptpn2^fl/fl^* controls (Fig. S4e-g), consistent with the possibility of decreased T cell exhaustion. AT3-OVA tumours in mice treated with PTPN2-deficent OT-1 CD8+ T cells started to re-emerge after 21 days, but survival was prolonged for as long as 86 days (**Fig. 2d**; Fig. S4h); by contrast control mice achieved the maximum ethically permissible tumour burden (200 mm^2^) by 25 days. Tumour re-emergence in this setting was accompanied by decreased OVA and MHC class I (*H2-k1*) gene expression, consistent with decreased antigen-presentation; tumour re-emergence was also accompanied by decreased PD-L1 (*Cd274*) gene expression (**Fig. 2e**), but this probably followed decreased MHC class I-mediated antigen presentation and thereby T cell recruitment and inflammation. Taken together these results are consistent with PTPN2 deficiency increasing the functional activity and attenuating the tolerisation of naïve CD8+ T cells to suppress tumour growth.

**Figure 2.**
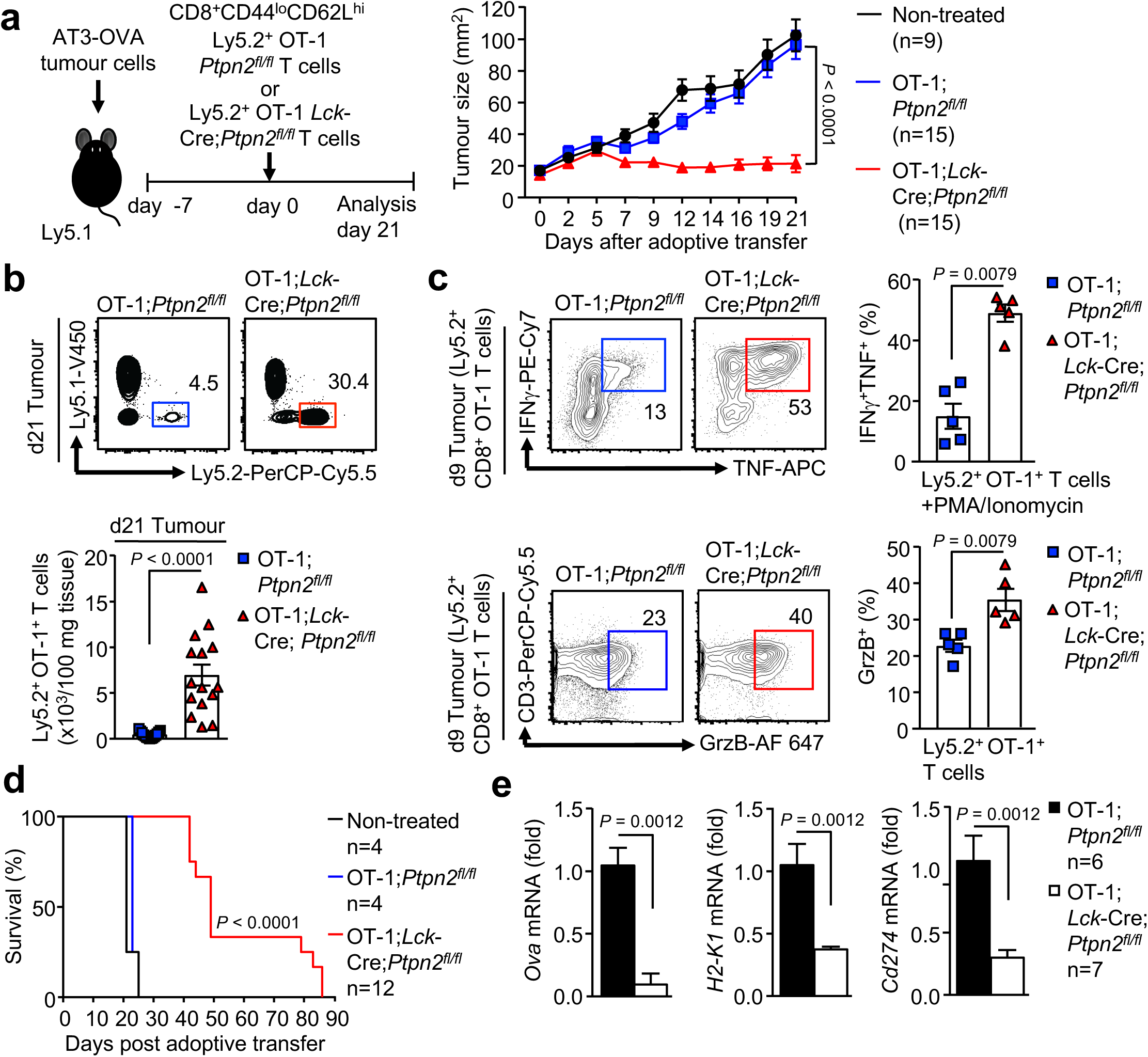
PTPN2-deletion enhances CD8+ T cell-mediated immunosurveillance. **a-d**) AT-3-OVA mammary tumour cells (1×10^6^) were injected into the fourth inguinal mammary fat pads of female Ly5.1^+^ mice. Seven days after tumour injection FACS-purified 2×10^6^ naïve CD8^+^CD44^lo^CD62L^hi^ lymph node T cells from Ly5.2^+^;OT-1;*Ptpn2^fl/fl^* versus Ly5.2^+^;OT-1;*Lck*-Cre;*Ptpn2^fl/fl^* mice were adoptively transferred into tumour-bearing Ly5.1 mice. Tumour-bearing Ly5.1 mice were monitored for **a**) tumour growth over 21 days and **d**) for survival over 86 days. **b**) After 21 days TILs were stained with fluorochrome-conjugated antibodies for Ly5.1 and Ly5.2, and Ly5.2^+^;OT-1;*Ptpn2^fl/fl^* and Ly5.2^+^;OT-1;*Lck*-Cre;*Ptpn2^fl/fl^* donor T cell numbers determined. **c**) After 9 days isolated TILs were stained for Ly5.1 and Ly5.2, as well as intracellular IFNγ, TNF and Granzyme B (GrzB) and the proportion of Ly5.2^+^IFNγ^+^TNF^+^ and Ly5.2^+^GrzB^+^ T cells determined. **e**) Gene expression in tumours from mice treated with Ly5.2^+^;OT-1;*Ptpn2^fl/fl^* T cells 21 days post-adoptive transfer versus those re-emerging in mice treated with Ly5.2^+^;OT-1;*Lck*-Cre;*Ptpn2^fl/fl^* T cells. Representative flow cytometry profiles and results (means ± SEM) from at least two independent experiments are shown. In (a) significance was determined using 2-way ANOVA Test and in (b, c, e,) significance determined using 2-tailed Mann-Whitney U Test. In (d) significance was determined using Log-rank (Mantel-Cox) test.

### PTPN2-deficiency enhances CAR T cell cytotoxicity

CAR T cells are autologous T cells engineered to express a transmembrane CAR specific for a defined tumour antigen that signals via canonical TCR signalling intermediates such as LCK ^23, 34^. CAR T cells targeting CD19 have especially been impressive in the treatment of ALL, with clinical response rates of up to 90% in pediatric B cell ALL patients ^20, 21^. However, therapeutic efficacies of CAR T cells in other malignancies, including solid tumours, have been relatively poor ^22, 23^. Given our findings on PTPN2 in T cell mediated immunosurveillance and anti-tumour immunity, we determined whether targeting PTPN2 might enhance the function of CAR T cells in solid tumours. In particular, we assessed the therapeutic efficacy of second-generation CAR T cells harboring the intracellular signalling domains of CD28 and CD3ζ and targeting the human orthologue of murine ErbB2/Neu, HER-2 ^35^. HER-2 is overexpressed in many solid tumours, including 20% of breast cancers, where it promotes tumour aggressiveness and metastasis ^36^.

First, we assessed the impact of PTPN2 deletion on CAR T cells *in vitro*. Consistent with PTPN2’s role in setting thresholds for TCR-instigated responses ^5, 10, 11^, we found that PTPN2-deficiency resulted in ten-fold lower concentrations of TCR crosslinking antibodies (α-CD3ε) being required for the maximal generation of CD8^+^ HER-2 CAR T cells *in vitro*, with the resulting CAR T cells being predominated by the effector/memory (CD44^hi^CD62L^lo^) subset (Fig. S5a). Next, we assessed the impact of PTPN2-deficiency on antigen-induced CAR T cell activation *in vitro*. PTPN2-deficient CD8^+^ HER-2 CAR T cells were more activated, as assessed by the expression of CD44, CD25, PD-1 and LAG-3, after overnight incubation with HER-2-expressing 24JK (24JK-HER-2) sarcoma cells, but importantly, not HER-2-negative 24JK control cells (Fig. S5b). Moreover, PTPN2-deficient CD8^+^ HER-2 CAR T cells exhibited increased antigen-specific cytotoxic capacity *in vitro* (Fig. S5c; Fig. S6a), as assessed by the increased intracellular expression of IFNγ (required for tumour eradication by CAR T cells *in vivo* ^37^), TNF and granzyme B upon challenge with 24JK-HER-2 cells but not 24JK cells. Moreover, both effector/memory and central memory (CD44^hi^CD62L^hi^) PTPN2-deficient CD8+ HER-2 CAR T cells were more effective at specifically killing 24JK-HER-2 cells but not 24JK cells *in vitro* (Fig. S6b). Taken together these results are consistent with PTPN2-deficiency enhancing not only the generation, but also the antigen-specific activation and cytotoxicity of CAR T cells *in vitro*.

### PTPN2-deficiency enhances LCK-dependent CAR T cell function

Next, we explored the mechanisms by which PTPN2-deficiency may influence CAR T cell activation and function. PTPN2 dephosphorylates and inactivates the SFK LCK to tune TCR signalling so that T cells can differentially respond to self versus non-self ^5, 9–11^. CAR T cells are reliant on canonical TCR signalling intermediates, including LCK for their activation and function ^34^. Accordingly, we assessed the influence of PTPN2 deficiency on the activation of LCK in CAR T cells by monitoring for the phosphorylation of Y394 (using antibodies specific for Y418-phosphorylated SFKs). PTPN2-deficiency significantly increased SFK Y418 phosphorylation in CD8+ HER-2 CAR T cells (Fig. S7a; **Fig. 3a**). We have shown previously that the enhanced TCR-induced T cell activation resulting from PTPN2 deficiency is accompanied by the increased expression of the interleukin (IL)-2 receptor chains CD25 (IL-2 receptor α chain) and CD122 (IL-2/15 receptor β chain) and IL-2 induced STAT-5 signalling ^5, 11^.

**Figure 3.**
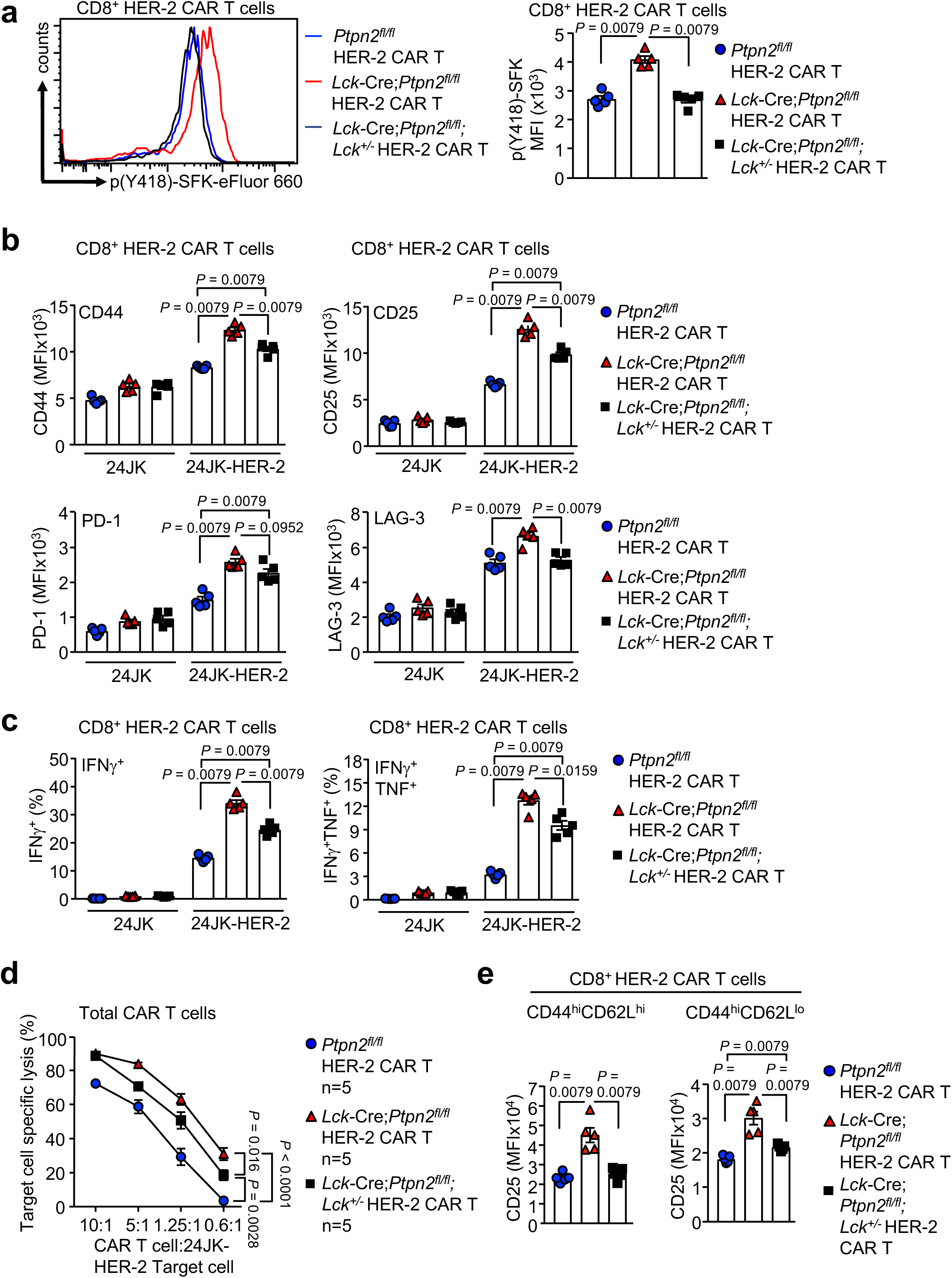
PTPN2-deletion enhances the LCK-dependent activation of CAR T cells. **a**) CD8^+^ HER-2 CAR T cells generated from *Ptpn2^fl/fl^* versus *Lck*-Cre;*Ptpn2^fl/fl^* versus *Lck*-Cre;*Ptpn2^fl/fl^*;*Lck^+/-^* splenocytes were stained for intracellular p(Y418)-SFK and p(Y418)-SFK MFIs were determined by flow cytometry. **b**) HER-2-specific *Ptpn2^fl/fl^* versus *Lck*-Cre;*Ptpn2^fl/fl^* versus *Lck*-Cre;*Ptpn2^fl/fl^*;*Lck^+/-^* CAR T cells were incubated with HER-2 expressing 24JK sarcoma cells (24JK-HER-2) or HER-2 negative 24JK sarcoma cells. Cells were stained for CD8, CD25, CD44, PD-1 and Lag-3 and CD44, CD25, PD-1 and LAG-3 MFIs were determined by flow cytometry. **c**) *Ptpn2^fl/fl^*, *Lck*-Cre;*Ptpn2^fl/fl^*, or *Lck*-Cre;*Ptpn2^fl/fl^*;*Lck^+/-^* HER-2 CAR T cells were incubated with 24JK-HER-2 or 24JK sarcoma cells. Cells were stained for CD8, intracellular IFNγ and TNF, and the proportion of CD8^+^IFNγ^+^ and CD8^+^IFNγ^+^TNF^+^ CAR T cells determined by flow cytometry. **d**) *Ptpn2^fl/fl^*, *Lck*-Cre;*Ptpn2^fl/fl^* or *Lck*-Cre;*Ptpn2^fl/fl^*;*Lck^+/-^* HER-2 CAR T cells were incubated with 5 mM CTV-labelled (CTV^bright^) 24JK-HER-2 and 0.5 mM CTV-labelled (CTV^dim^) 24JK sarcoma cells. Antigen-specific target cell lysis was monitored for the depletion of CTV^bright^ 24JK-HER-2 cells by flow cytometry. **e**) HER-2-specific *Ptpn2^fl/fl^*, *Lck*-Cre;*Ptpn2^fl/fl^* and *Lck*-Cre;*Ptpn2^fl/fl^*;*Lck^+/-^* CAR T cells were incubated with plate-bound α-CD3 and stained for CD8, CD44, CD62L and CD25 and CD25 MFIs on CD8^+^CD44^hi^CD62L^lo^ versus CD8^+^CD44^hi^CD62L^hi^CAR T cells determined by flow cytometry. Representative histograms and results (means ± SEM) from at least two independent experiments are shown. In (a-c, e) significance determined using 2-tailed Mann-Whitney U Test. In (d) significance was determined using 2-way ANOVA Test.

Consistent with this we found that CD25, CD122 and CD132 (common γ chain shared by IL-2 and IL-15) receptor levels were elevated in activated PTPN2-deficient CAR T cells (Fig. S7b; **Fig. 3b**) and this was accompanied by increased basal and IL-2/15-induced STAT-5 Y694 phosphorylation (Fig. S7c-d). To explore the extent to which the increased SFK signalling may contribute to this and the enhanced CAR T cell activation and function we crossed *Lck*-Cre;*Ptpn2^fl/fl^* mice onto the *Lck^+/–^* background^9^ so that total LCK would be reduced by 50% and LCK signalling may more closely approximate that in *Ptpn2^fl/fl^* controls. Consistent with this we found that SFK Y418 phosphorylation in *Lck*-Cre;*Ptpn2^fl/fl^*;*Lck^+/–^* CD8+ HER-2 CAR T cells was reduced to that in *Ptpn2^fl/fl^* controls (**Fig. 3a**). Strikingly, *Lck* heterozygosity attenuated the enhanced antigen-specific HER-2 CAR T cell activation (as monitored by CD44, CD25, PD-1 and LAG-3 levels; **Fig. 3b**) and cytotoxic potential (as measured by the antigen-induced expression of IFNγ and TNF; **Fig. 3c**) and the enhanced capacity of PTPN2-deficient CAR T cells to specifically kill HER-2-expressing tumour cells (**Fig. 3d**). In addition, *Lck* heterozygosity corrected the enhanced IL-2/15 receptor levels (**Fig. 3e**; Fig. S7e) and partially corrected the enhanced IL-2/15-induced STAT-5 signalling (Fig. S7f). The persistent increased STAT-5 signalling despite correcting IL-2/15 receptor levels is consistent with previous studies showing that STAT-5 can also serve as direct a *bona-fide* substrate of PTPN2 and that PTPN2-deficiency promotes cytokine-induced STAT-5 signalling in thymocytes/T cells ^5, 6, 9, 10, 38, 39^. Irrespective, these results are consistent with PTPN2 deficiency enhancing the antigen-specific activation and function of CAR T cells through the promotion of LCK signalling.

### PTPN2-deficient CAR T cells eradicate solid tumours

To explore the therapeutic efficacy of PTPN2-deficient HER-2-targeting CAR T cells *in vivo* we adoptively transferred a single dose (6 × 10^6^) of purified *Ptpn2^fl/fl^* versus *Lck*-Cre;*Ptpn2^fl/fl^* central memory CD8+ HER-2 CAR T cells into sub-lethally irradiated syngeneic recipients bearing established orthotopic tumours arising from the injection of HER-2-expressing E0771 (HER-2-E0771) breast cancer cells (**Fig. 4**). We adoptively transferred central memory CAR T cells as these cells engraft better and elicit persistent anti-tumour responses ^40^. In addition, as lymphodepletion prior to T cell infusion is used routinely in the clinic to facilitate T cell expansion and enhance efficacy, ^41^ we immunodepleted mice by sublethal irradiation (400 cGy) as is routine in murine CAR T cell studies ^42^. Notably, HER-2-E0771 cells were grafted into HER-2 transgenic (TG) mice, where HER-2 expression was driven by the whey acidic protein (WAP) promoter that induces expression in the cerebellum and the lactating mammary gland ^43^, so that HER-2-expressing orthotopic tumours would be regarded as self and host anti-tumour immunity repressed. Previous studies have shown that effective tumour killing and eradication by CD8+ CAR T cells is reliant on the presence of CD4+ CAR T cells ^37^. Strikingly, we found that although PTPN2-expressing central memory CD8+ HER-2 CAR T cells modestly suppressed HER-2-E0771 mammary tumour growth, PTPN2-deficient CD8+ HER-2 CAR T cells eradicated tumours (**Fig. 4a**). The ability of PTPN2-deficient CAR T cells to suppress tumour growth and eradicate tumours was reliant on the enhanced activation of LCK, as this was significantly, albeit not completely abrogated by *Lck* heterozygosity (**Fig. 4b**) that corrected the enhanced LCK activation in *Lck*-Cre;*Ptpn2^fl/fl^* CAR T cells *in vitro* (**Fig. 3a**). The repression of tumour growth by PTPN2-deficient CD8+ HER-2 CAR T cells occurred despite tumours harboring an immunosuppressive microenvironment ^44, 45^, with increased immunosuppressive myeloid-derived suppressor cells (MDSCs) (**Fig. 4c**) and T_regs_ (**Fig. 4d**) and the increased expression of immunosuppressive cytokines, including transforming growth factor β (*Tgfb*) and IL-10 (*Il10*) (**Fig. 4e**), when compared to normal mammary tissue. PTPN2-deficient CAR T cells markedly suppressed tumour growth, even when the tumours were allowed to grow to one quarter (50 mm^2^) of the maximal ethically permissible mammary tumour burden prior to CAR T cell therapy (**Fig. 4f**). Moreover, PTPN2-deficient CAR T cells were effective in repressing tumour growth even without the co-administration of IL-2 that is used routinely in rodent pre-clinical models to promote CAR T cell expansion, but is not used in the clinic (**Fig. 4g**). Taken together these results demonstrate that PTPN2-deficiency promotes the LCK-dependent activation of CAR T cells and overcomes the immunosuppressive tumour microenvironment to eradicate solid tumours *in vivo*.

**Figure 4.**
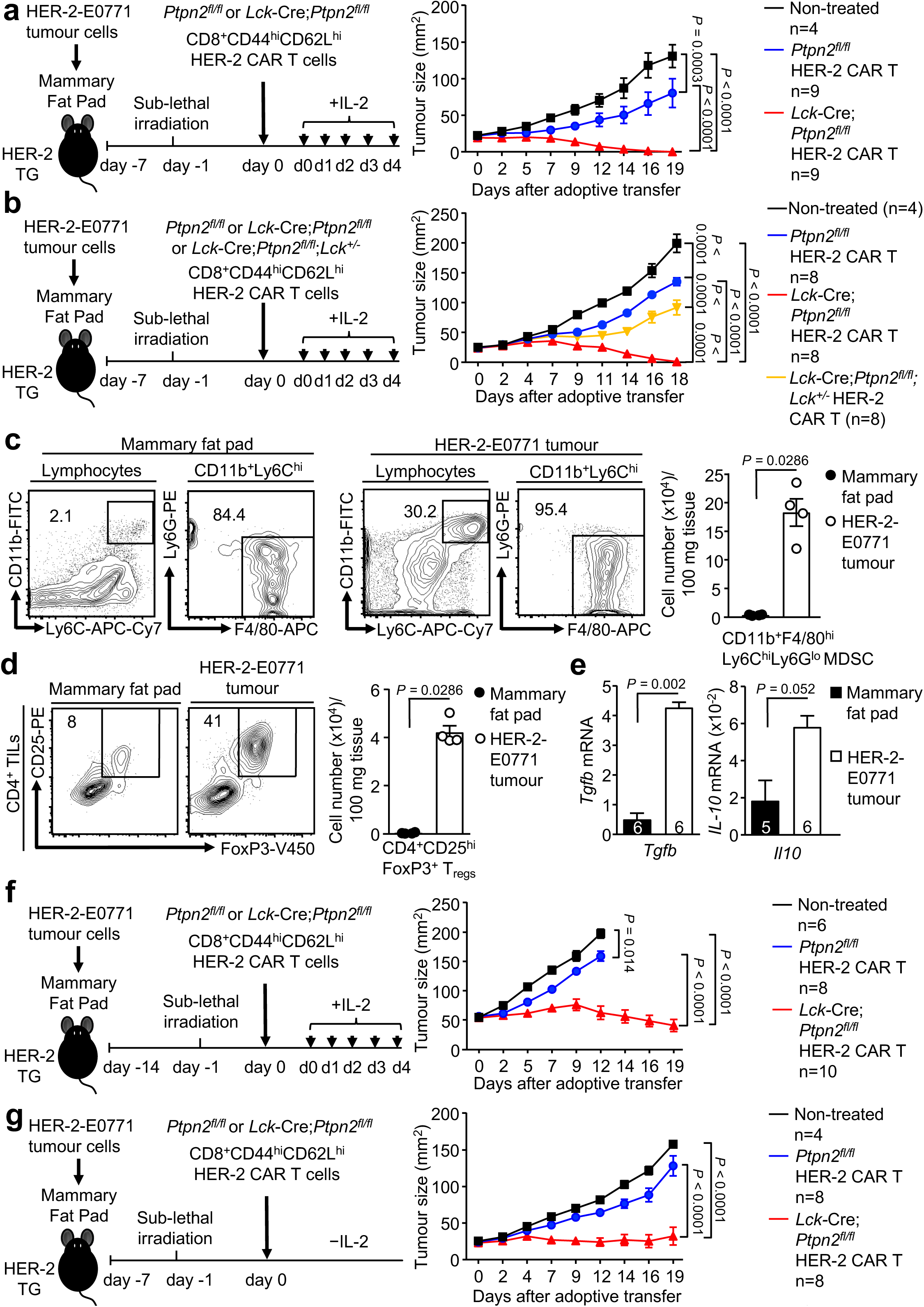
PTPN2-deletion enhances CAR T cell efficacy in vivo. **a, b, f, g**) HER-2-E0771 mammary tumour cells (2×10^5^) were injected into the fourth inguinal mammary fat pads of female HER-2 TG mice. **a, b, g**) Seven days or **f**) 14 days after tumour injection HER-2 TG mice received total body irradiation (4Gy) followed by the adoptive transfer of 6×10^6^ FACS-purified CD8^+^CD44^hi^CD62L^hi^ central memory HER-2 CAR T cells generated from *Ptpn2^fl/fl^* versus *Lck*-Cre;*Ptpn2^fl/fl^* splenocytes. Mice were injected with **a, b, f**) IL-2 (50,000 IU/day) or **g**) saline on days 0-4 after adoptive CAR T cell transfer and monitored for tumour growth. **c-d**) TILs isolated from HER-2-E0771 mammary tumour were stained for **c**) CD11b, F4/80, Ly6C and Ly6G or **d**) CD4, CD25 and intracellular FoxP3 and the number of **c**) CD11b^+^ F4/80^hi^Ly6C^hi^Ly6G^lo^ myeloid derived suppressor cells (MDSC) and **d**) CD4^+^CD25^+^FoxP3^+^ regulatory T cells (T_regs_) determined by flow cytometry. **e**) *Tgfb* and *IL-10* mRNA levels in HER-2-E0771 tumours were assessed by quantitative real time PCR. Representative flow cytometry profiles and results (means ± SEM) from three independent experiments are shown. In (a, b, f, g) significance was determined using 2-way ANOVA Test. In (c-e) significance was determined using 2-tailed Mann-Whitney U Test.

Strikingly the repression of tumour growth by PTPN2-deficient HER-2 CAR T cells persisted long after HER-2 tumours had been eradicated, so that approximately half of all mice were alive for longer than 200 days with the remaining half succumbing to tumours (**Fig. 5a**). Moreover, the tumours that did re-emerge lacked HER-2 as assessed by immunohistochemistry (**Fig. 5b**) and quantitative real time PCR (**Fig. 5c**).

**Figure 5.**
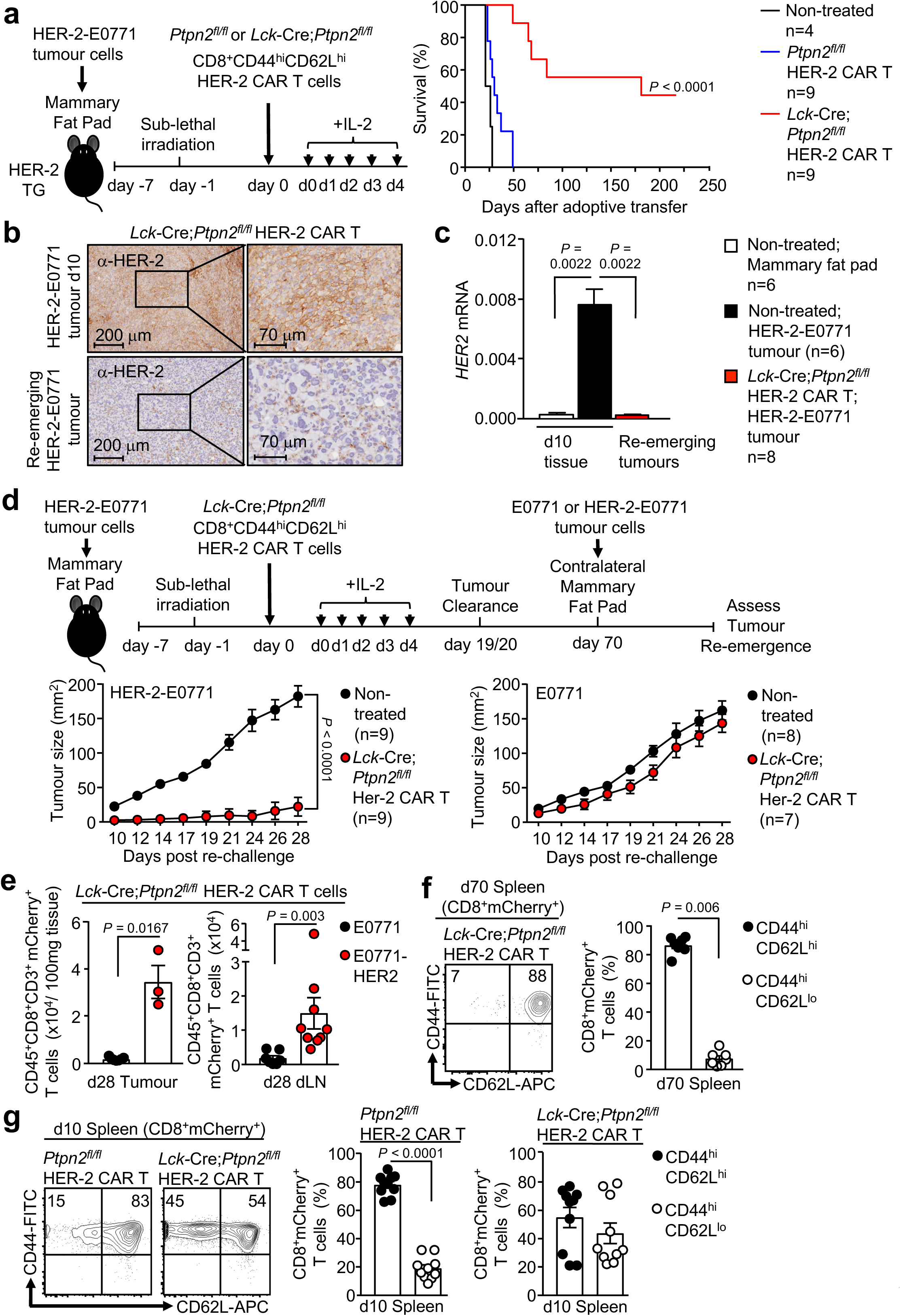
PTPN2-deficiency prevents the re-emergence of tumours. **a**) HER-2-E0771 mammary tumour cells (2×10^5^) were injected into the fourth inguinal mammary fat pads of female HER-2 TG mice. Seven days after tumour injection HER-2 TG mice received total body irradiation (4 Gy) followed by the adoptive transfer of 6×10^6^ FACS-purified *Lck*-Cre;*Ptpn2^fl/fl^* CD8^+^CD44^hi^CD62L^hi^ central memory HER-2 CAR T cells. Mice were injected with IL-2 (50,000 IU/day) on days 0-4 after adoptive CAR T cell transfer and monitored for survival. **b**) HER-2-E0771 tumours at day 10 post adoptive CAR T cell transfer or from HER-2-E0771 tumours that had re-emerged after being cleared by *Lck*-Cre;*Ptpn2^fl/fl^* CD8^+^ HER-2 CAR T cells were analyzed for HER-2 expression by immunohistochemistry. **c**) Normal mammary tissue or HER-2-E0771 tumours at day 10 post-adoptive CAR T cell transfer, or from those had re-emerged after being cleared by *Lck*-Cre;*Ptpn2^fl/fl^* CD8^+^ HER-2 CAR T cells were analysed for HER-2 expression by quantitative PCR. **d**) HER-2-E0771 versus HER-2 negative E0771 breast cancer cells (2×10^5^) were injected into the contralateral fourth inguinal mammary fat pads of female HER-2 TG mice 50 days after adoptive CAR T cell transfer and mice were monitored for tumour growth. **e**) At day 28 the number of CD45^+^CD3^+^CD8^+^ mCherry^+^ CAR T cells in HER-2-E0771 versus E0771 tumours and dLNs were determined by flow cytometry. **f-g**) *Ptpn2^fl/fl^* versus *Lck*-Cre;*Ptpn2^fl/fl^* HER-2 CAR T cells isolated from the spleens of HER-2 TG mice at **f**) 70 days and **g**) 10 days post adoptive transfer were stained for CD8, CD44 and CD62L and analysed by flow cytometry. Representative flow cytometry profiles and results (means ± SEM) from three independent experiments are shown. In (a) significance was determined using Log-rank (Mantel-Cox) test. In (d) significance was determined using 2-way ANOVA Test. In (e-g) significance was determined using 2-tailed Mann-Whitney U Test.

These results are consistent with PTPN2-deficient HER-2 CAR T cells completely eliminating HER-2 expressing tumours and eliciting a selective pressure so that any re-emerging tumours downregulate HER-2. To explore this further and the influence of PTPN2-deficiency on CAR T cell memory, we re-implanted HER-2+ tumour cells into control HER-2 transgenic mice, or those mice in which HER-2+ tumours had been previously cleared by PTPN2-deficient HER-2 CAR T cells (Fig. 5d-e). In those mice in which tumours had previously been cleared, splenic PTPN2-deficient CAR T cells had a central memory phenotype (CD44^hi^CD62L^hi^) rather than the mixed central and effector/memory (CD44^hi^CD62L^lo^) phenotypes otherwise present on day 10 post adoptive transfer (Fig. 5f-g), consistent with PTPN2 deficiency promoting CAR T cell memory. To assess systemic anti-tumour immunity, HER-2+ tumours cells were re-implanted into the previously tumour-free contralateral mammary fat pads. As a control, we also monitored the growth of HER-2-negative E0771 tumour cells. We found that the growth of HER-2+ tumours in the previously tumour-free contralateral mammary fat pads was markedly repressed or completely prevented (**Fig. 5d**). The repression of tumour growth was accompanied by the increased infiltration of HER-2 CAR T cells (**Fig. 5e**). By contrast, the growth of HER-2-negative mammary E0771 tumours was not affected (**Fig. 5d**). These results are consistent with PTPN2-deficiency in HER-2 CAR T cells promoting CAR T cell memory and recall to prevent the re-emergence of HER-2+ tumours, including those that may arise at distant metastatic sites.

### PTPN2-deficiency promotes CAR T cell homing

The clinical benefit of adoptive T cell therapy is highly reliant on the ability of T cells including CAR T cells to home efficiently into the target tissue ^46, 47^. We found that the repression of tumour growth by PTPN2-deficient CD8+ HER-2 CAR T cells was accompanied by a marked increase in CD45^+^CD8^+^CD3^+^mCherry^+^ CAR T cell abundance in tumours and the corresponding draining lymph nodes (Fig. 6a). In part, the increased CAR T cell abundance may reflect the expansion of CAR T cells after they engage tumour antigen, as PTPN2-deficiency increased the antigen-specific proliferation of HER-2 CAR T cell *in vitro* (Fig. 6b; Fig. S8a). However, the increased CAR T cell abundance may also reflect an increase in CAR T cell homing and infiltration as CD3ε+ lymphocytes accumulated within the tumour and at the tumour/stromal interface at 10 days post adoptive transfer (Fig. S8b). Previous studies have correlated the accumulation of TILs in tumours with the expression of the chemokine receptor CXCR3 (receptor for CXCL9, CXCL10 and CXCL11) ^46, 47^. We found that PTPN2-deficient HER-2 CAR T cells expressed higher cell surface levels of the chemokine receptor CXCR3. By contrast, cell surface levels of other chemokine receptors, including CXCR5, CCR7 and CCR5 (receptor for CCL3, CCL4 and CCL5), were not altered (Fig. 6c-d; Fig. S8c). The ligands for CXCR3 are increased in many tumours and associated with the intralesional accumulation of TILs and improved outcome ^46, 47^ and *Cxcl9* and *Cxcl10* (*Cxcl11* is not expressed in C57BL/6 mice ^48^) were elevated in the HER-2-E0771 tumours analysed at 10 days after implantation (Fig. 6e). Moreover, consistent with the potential for increased homing, we found that PTPN2-deficient CXCR3^hi^ CAR T cells accumulated in HER-2-E0771 tumours within 3 days of adoptive transfer (Fig. 6f), prior to any effects on tumour burden (**Fig. 4a**).

**Figure 6.**
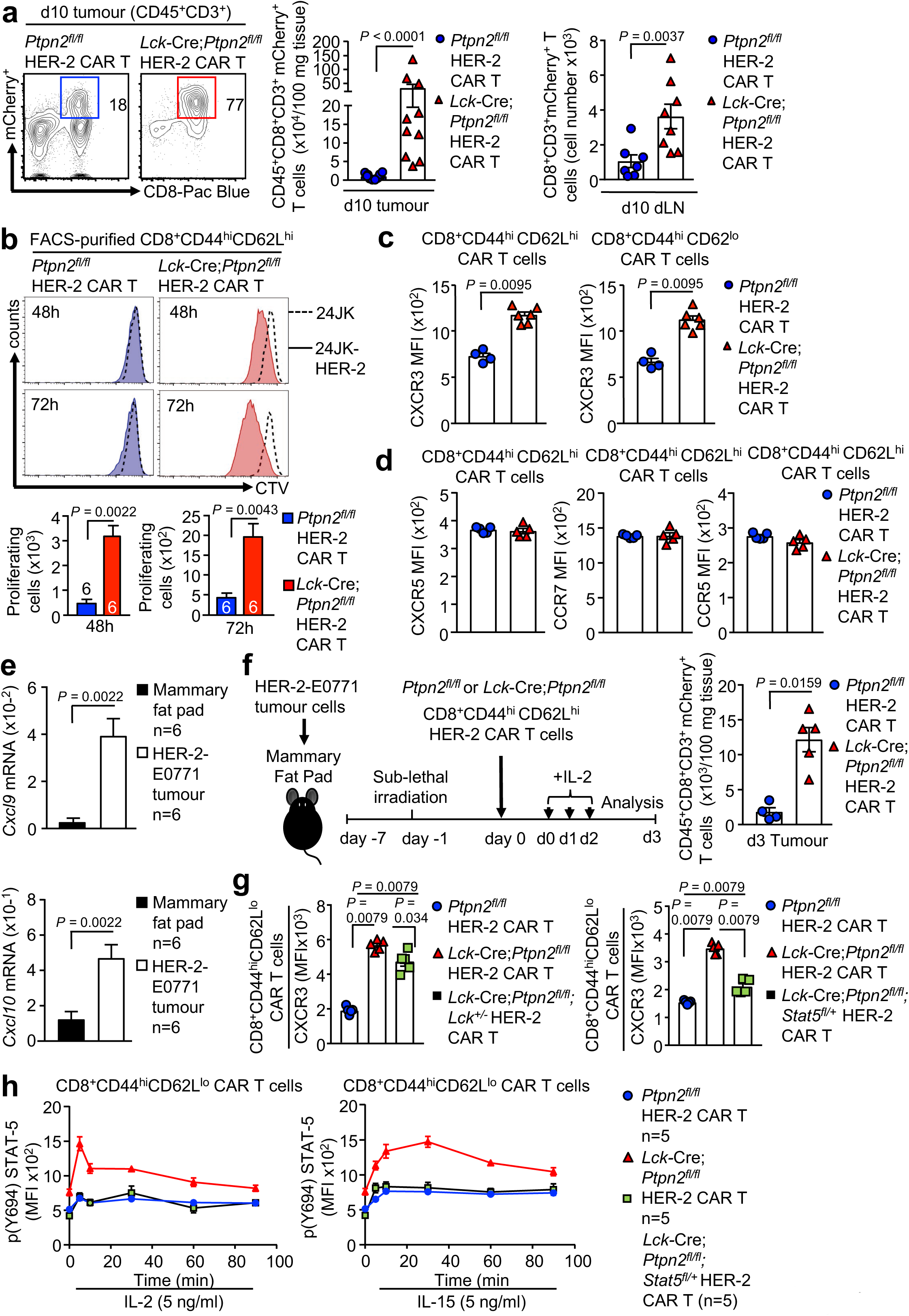
PTPN2-deficiency enhances CXCR3 expression and promotes CAR T cell homing. HER-2-E0771 mammary tumour cells (2×10^5^) were injected into the fourth inguinal mammary fat pads of female HER-2 TG mice. Seven days after tumour injection HER-2 TG mice received total body irradiation (4 Gy) followed by the adoptive transfer of 6×10^6^ FACS-purified CD8^+^CD62L^hi^CD44^hi^ central memory HER-2 CAR T cells generated from *Ptpn2^fl/fl^* or *Lck*-Cre;*Ptpn2^fl/fl^* splenocytes. Mice were injected with IL-2 (50,000 IU/day) on days 0-4 after adoptive CAR T cell transfer. **a**) Lymphocytes were isolated from the tumours and dLN at day 10 and **f**) at day 3 post adoptive transfer and stained for CD45, CD3, CD8 and mCherry^+^CD45^+^CD3^+^CD8^+^ CAR T cell numbers were determined by flow cytometry. **b**) CTV-labelled *Ptpn2^fl/fl^* versus *Lck*-Cre;*Ptpn2^fl/fl^* CD8^+^ HER-2 CAR T cells were incubated with 24JK-HER-2 or 24JK sarcoma cells and CTV dilution assessed by flow cytometry to monitor proliferation. **c-d**) HER-2 CAR T cells generated from *Ptpn2^fl/fl^* versus *Lck*-Cre;*Ptpn2^fl/fl^* splenocytes were stained for CD8, CD62L, CD44 and CXCR3, CXCR5, CCR7, CCR5 and **c**) CXCR3 MFIs on CD8^+^CD44^hi^CD62L^lo^ and CD8^+^CD44^hi^CD62L^hi^ CAR T cells and **d**) CXCR5, CCR7, CCR5 MFIs on CD8^+^CD44^hi^CD62L^lo^ CAR T cells were determined by flow cytometry. **e**) *Cxcl9* and *Cxcl10* mRNA levels in HER-2-E0771 tumours were assessed by quantitative real time PCR. **g**) *Ptpn2^fl/fl^* versus *Lck*-Cre;*Ptpn2^fl/fl^*, *Lck*-Cre;*Ptpn2^fl/fl^*;*Lck^+/-^* or *Lck*-Cre;*Ptpn2^fl/fl^*;*Stat5^fl/+^* HER-2 CAR T cells were incubated with plate-bound α-CD3 and stained for CD8, CD62L, CD44 and CXCR3 and CXCR3 MFIs on CD8^+^CD44^hi^CD62L^lo^ CAR T cells determined by flow cytometry. **h**) *Ptpn2^fl/fl^* versus *Lck*-Cre;*Ptpn2^fl/fl^* versus *Lck*-Cre;*Ptpn2^fl/fl^*;*Stat5^fl/+^* HER-2 CAR T cells were incubated with plate-bound α-CD3 and stimulated with murine recombinant IL-2 and IL-15 for the indicated time points. Cells were stained for CD8, CD62L, CD44 and intracellular p(Y694)-STAT5 MFIs in CD8^+^CD44^hi^CD62L^lo^ were determined by flow cytometry. Representative flow cytometry profiles and results (means ± SEM) from two independent experiments are shown. In (a, b, c, e, f, g) significance was determined using 2-tailed Mann-Whitney U Test.

To explore whether the increased cell surface CXCR3 might contribute to the increased homing and anti-tumour activity of PTPN2-deficient CAR T cells, we sought to correct the increased CXCR3 expression. CXCR3 is not detected in naïve T cells but is abundant in CD4+ TH1 cells and CD8+ cytotoxic T lymphocytes (CTLs). When CD8+ T cells are activated by a strong TCR stimulus and subsequently stimulated with IL-2 they undergo differentiation into effectors and acquire CTL activity characterized by IFNγ and granzyme B expression ^49, 50^. Our previous studies have shown that PTPN2-deficiency enhances the IL-2-induced generation of effectors from TCR crosslinked and activated T cells ^11^. Given that CAR T cell generation is reliant on TCR crosslinking (α-CD3ε/α-CD8) and stimulation with IL-2 and IL-7, we determined whether the enhanced LCK activation and increased downstream IL-2 receptor and STAT-5 signalling in PTPN2-deficient CAR T cells might be responsible for the increased CXCR3 expression. To assess this we took advantage of PTPN2-deficient CD8+ CAR T cells that were heterozygous for *Lck* (*Lck*-Cre;*Ptpn2^fl/fl^*;*Lck^+/–^*). Since IL-2-induced STAT-5 signalling was only partially corrected in *Lck*-Cre;*Ptpn2^fl/fl^*;*Lck^+/–^* CAR T cells (Fig. S7f) we also crossed the *Lck*-Cre;*Ptpn2^fl/fl^* mice onto the *Stat5^fl/+^*background so that we could independently correct the increased STAT-5 signalling (Fig. 6g-h). Cell surface CXCR3 levels were significantly albeit modestly reduced in *Lck* heterozygous CAR T cells (Fig. 6g). By contrast, *Stat5* heterozygosity completely corrected the enhanced IL-2 and IL-15-induced STAT-5 signalling and almost completely corrected the increased CXCR3 (Fig. 6g-h).

Although CXCR3 is not transcribed by STAT-5, there is evidence that STAT-5 can drive the expression of the transcription factor T-bet (T-box transcription factors T-box expressed in T cells) ^51^ which together with Eomes (eomesodermin) dictates CD8+ T cell differentiation and function. Indeed T-bet can drive the expression of CXCR3 ^52^ and has been shown to be important for the infiltration and anti-tumour activity of cytotoxic CD8+ T cells ^53^. Consistent with this we found that PTPN2-deficiency in CAR T cells was associated with increased intracellular T-bet expression that was corrected by *Stat5* heterozygosity (Fig. S8d-e). Strikingly, *Stat5* but not *Lck* heterozygosity prevented the increased homing of PTPN2-deficient CAR T cells evident at 3 days post adoptive transfer (**Fig. 7a**) and largely, albeit not completely, attenuated the ability of PTPN2-deficent CAR T cells to suppress the growth solid tumours (**Fig. 7b**). The repression of CAR T cell infiltration was also evident in resected tumours at day 16 with *Lck*-Cre;*Ptpn2^fl/fl^*;*Stat5^fl/+^* CAR T cell cytotoxicity markers (TNF, IFNγ; induced by PMA/ionomycin *ex vivo*) being reduced to those in *Ptpn2^fl/fl^*control CAR T cells (**Fig. 7c**). Therefore, the promotion of STAT-5 signalling might not only promote CXCR3 expression and the homing of CAR T cells to CXCL9/10-expressing tumours, but also contribute to the acquisition of CTL activity probably through the induction of T-bet and thereby CAR T cell function.

**Figure 7.**
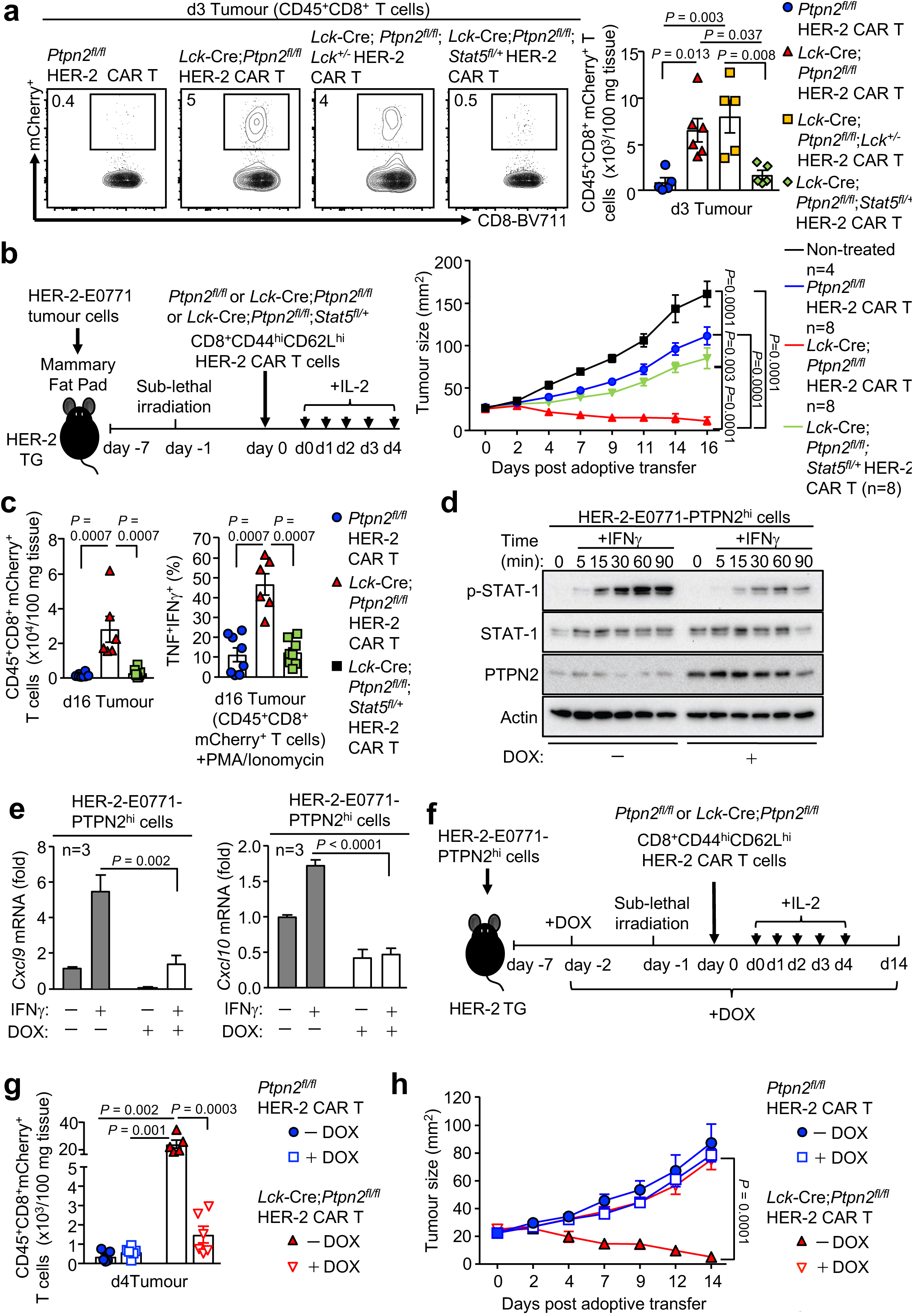
PTPN2-deficiency enhances CAR T cell efficacy in vivo by promoting STAT5-mediating homing to CXCL9/10-expressing tumours. **a-c**) HER-2-E0771 mammary tumours cells (2×10^5^) were injected into the fourth inguinal mammary fat pads of female HER-2 TG mice. Seven days after tumour injection HER-2 TG mice received total body irradiation (4 Gy) followed by the adoptive transfer of 6×10^6^ FACS-purified CD8^+^CD62L^hi^CD44^hi^ central memory HER-2 CAR T cells generated from *Ptpn2^fl/fl^*, *Lck*-Cre;*Ptpn2^fl/fl^*, *Lck*-Cre;*Ptpn2^fl/fl^*;*Stat5^fl/+^* or *Lck*-Cre;*Ptpn2^fl/fl^*;*Lck*^+/-^ splenocytes. Mice were injected with IL-2 (50,000 IU/day) on days 0-4 after adoptive CAR T cell transfer and monitored for tumour growth. **a, c**) Lymphocytes were isolated from the tumours on **a**) days 3 or **c**) day 16 post adoptive transfer and stained for CD45 and CD8 and mCherry^+^CD45^+^ CD8^+^ CAR T cell numbers determined by flow cytometry. In (c) TILs were stained for intracellular IFNγ and TNF after PMA/Ionomycin treatment. **d**) HER-2-E0771 cells generated to inducibly overexpress PTPN2 in response to doxycycline (E0771-HER-2-PTPN2^hi^) were pre-incubated (24 h) with vehicle or doxycycline (DOX) subsequently stimulated with IFNγ for the indicated times. STAT-1 Y701 phosphorylation (p-STAT-1) and PTPN2 levels were assessed by immunoblotting. **e**) *Cxcl9* and *Cxcl10* mRNA levels in vehicle versus DOX-treated and IFNγ-stimulated HER-2-E0771 cells were assessed by quantitative real time PCR. **f-h**) E0771-HER-2-PTPN2^hi^ mammary tumour cells (2×10^5^) were injected into the fourth inguinal mammary fat pads of female HER-2 TG mice. Five days after tumour injection mice were administered vehicle or DOX in drinking water followed by irradiation (4 Gy) on day 7 and the adoptive transfer of 6×10^6^ FACS-purified central memory *Ptpn2^fl/fl^* versus *Lck*-Cre;*Ptpn2^fl/fl^* HER-2 CAR T cells. Mice were then injected with IL-2 (50,000 IU/day) on days 0-4 post adoptive CAR T cell transfer and tumour growth was monitored. In (g) CD45^+^CD8^+^ TILs were quantified by flow cytometry at day 4 post adoptive transfer. Representative flow cytometry profiles and results (means ± SEM) from two independent experiments are shown. In (a, e) significance was determined using 1-way ANOVA Test. In (b, g, h) significance was determined using 2-way ANOVA Test. In (c) significance was determined using 2-tailed Mann-Whitney U Test.

To complement these findings and further explore the extent to which the CXCR3/CXCL9/10/11 axis might contribute to the increased homing and efficacy of PTPN2-deficient CAR T cells we sought to repress the *Cxcl9/10* expression in HER-2-E0771 mammary tumours and assess the impact on the homing and function of PTPN2-deficient CAR T cells. As *Cxcl9/10* are transcriptional targets of STAT-1 and PTPN2 dephosphorylates STAT-1 to repress IFNγ-induced STAT-1-mediated transcription^7, 38, 54, 55^, we generated HER-2-E0771 cells in which PTPN2 could be inducibly overexpressed in response to doxycycline (**Fig. 7d**). The inducible overexpression of PTPN2 not only repressed IFNγ-induced p-STAT-1 (**Fig. 7d**) and *Cxcl9/10* expression (**Fig. 7e**), but most importantly, also the recruitment of PTPN2-deficient CAR T cells to HER-2-E0771 mammary tumours *in vivo* (Fig. 7f-h). Importantly, although PTPN2-deficient CAR T cells remained more effective than controls, the doxycycline-inducible overexpression of PTPN2 in HER-2-E0771 cells and the decreased CAR T cell recruitment attenuated the repression of tumour growth and prevented the eradication of tumours (Fig. 7g-h). Taken together our findings are consistent with the efficacy of PTPN2-deficient CAR T cells in solid tumours being attributed to i) the increased LCK-dependent activation of CAR T cells after antigen engagement, ii) the LCK and STAT-5-dependent acquisition of CTL activity and iii) the increased STAT-5 mediated and CXCR3-dependent homing of PTPN2-deficient CAR T cells to CXCL9/10-expressing tumours.

### PTPN2-deficient CAR T cells do not promote morbidity

A potential complication of enhancing the function of CAR T cells is the development of systemic inflammation and autoimmunity ^23, 56^. We found that the eradication of tumours by PTPN2-deficient HER-2 CAR T cells was not accompanied by systemic T cell activation and inflammation, as assessed by the unaltered number and activation of T cells in lymphoid organs at 21 days post-transfer (Fig. S9a) and the unaltered circulating pro-inflammatory cytokines (Fig. S9b) and lymphocytic infiltrates in non-lymphoid tissues, including in the contralateral tumour-negative mammary glands, lungs and livers (Fig. S9c; data not shown). To further explore any impact of PTPN2-deficiency on the development of inflammatory disease, we increased the number of adoptively transferred CAR T cells from 6 × 10^6^ to 20 x 10^6^ and monitored for inflammation and autoimmunity. The increased CAR T cell numbers resulted in a more profound repression of tumour growth irrespective of PTPN2 status but only PTPN2-deficient CAR T cells eradicated tumours (Fig. S9d). PTPN2-deficiency did not exacerbate systemic inflammation, as assessed by monitoring for lymphocytic infiltrates in non-lymphoid tissues, including the contralateral mammary glands (data not shown), lungs and livers (Fig. S9e) or for circulating IL-6, IFNγ, TNF and IL-10 over time (Fig. S10a). Although PTPN2-deficient CAR T cells increased core temperature as early as 5 days post adoptive transfer (Fig. S10b), this is to be expected for a developing immune response and this did not persist after tumours were cleared and did not affect body weight (Fig. S10b-c). In addition, the increased PTPN2-deficient CAR T cells did not result in autoimmunity as reflected by the absence of circulating anti-nuclear antibodies (Fig. S10d) and the lack of any overt tissue damage, including liver damage, as assessed by measuring the liver enzymes alanine transaminase (ALT) and aspartate transaminase (AST) in serum (Fig. S10e). Therefore, PTPN2-deficient CAR T cells eradicate tumours without promoting systemic inflammation and immunopathologies.

Another potential complication of CAR T cell therapy is ‘on-target off-tumour’ toxicities ^23, 56^. Previous studies have shown that the WAP promoter in the HER-2 TG mice drives HER-2 in the lactating mammary gland and the cerebellum ^43^. Although PTPN2-deficient CAR T cells were not evident in the contralateral tumour-negative mammary glands, this is not unexpected as we used virgin mice in our studies. By contrast the WAP promoter ^43^ drives HER-2 expression in the cerebellum and we could detect HER-2 throughout the cerebellum (Fig. S11a) to similar levels seen in HER-2-E0771 mammary tumours (Fig. S11b). Although we detected some CD3^+^ mCherry^+^ CAR T cells surrounding the crus 1 of the ansiform lobule of the cerebelum they were not detected in other regions (Fig. S11c) and was there no evidence for overt cerebellar tissue damage (Fig. S11d), as assessed on day 10 post-adoptive transfer when PTPN2-deficient CAR T cells were activated (Fig. S11e). Consistent with this, PTPN2-deficient CAR T cells did not result in overt morbidity up to 50 days post-transfer and did not affect the cerebellar control of neuromotor function, as assessed in rotarod tests (Fig. S11f), even in those mice were a greater number of CAR T cells were adoptively transferred (data not shown). In part, the lack of toxicity may be due to CXCR3-expressing CAR T cells being unable to efficiently home and infiltrate into the cerebellum, as the cerebellum expressed negligible levels of *Cxcl9* and *Cxcl10* when compared to the HER-2-E0771 tumours (Fig. S11g). Therefore, targeting PTPN2 may not only enhance the antigen-specific activation and function of CAR T cells, but also limit ‘on-target off-tumour’ toxicities by driving the homing of CXCR3-expressing CAR T cells to CXCL9/10/11-expressing tumours. Taken together, our results demonstrate that PTPN2 deletion in murine CD8+ CAR T cells dramatically enhances their recruitment into the tumour site, as well as their antigen-specific activation and ability to overcome the immunosuppressive tumour microenvironment to effectively suppress the growth of solid tumours without promoting overt morbidity.

### PTPN2 inhibition enhances the antigen-specific activation of human CAR T cells

To explore whether targeting PTPN2 in human T cells and CAR T cells might similarly promote their activation and function we took advantage of a highly specific PTPN2 active site inhibitor, compound 8 ^58, 59^. Treatment of murine CD8+ HER-2 CAR T cells with compound 8 *in vitro* increased their antigen-specific cytotoxic potential to levels seen in *Lck*-Cre;*Ptpn2^fl/fl^* HER-2 CAR T cells, but had no additional effect on *Lck*-Cre;*Ptpn2^fl/fl^* HER-2 CAR T cells, consistent with the inhibitor acting specifically to inhibit PTPN2 and thereby activate CAR T cells upon engagement with specific tumour antigen (Fig. S12a). Importantly, we found that treatment of human peripheral blood mononuclear cells with compound 8 enhanced their activation (as assessed by the activation marker CD69) and the expansion of CD8^+^CCR7^+^CD45RA^+^CD69^+^ T cells in response to TCR ligation (α-CD3ε) (Fig. S12b). To assess whether targeting of PTPN2 might enhance the cytotoxic potential of human CAR T cells we took advantage of human CAR T cells targeting the Lewis Y (LY) antigen ^60, 61^ that is overexpressed in many human cancers, including 80% of lung adenocarcinomas, 25% of ovarian carcinomas and 25% of colorectal adenocarcinomas (**Fig. 8a**). Treatment of human LY CAR T cells with compound 8 significantly enhanced their cytotoxic potential, as assessed by the expression of IFNγ and TNF, in response to CAR crosslinking (with α-LY) or engagement with LY-expressing human ovarian (OVCAR-3) carcinoma cells, but not breast cancer cells (MDA-MB-231) that do not express LY (**Fig. 8a**). Taken together these results are consistent with PTPN2 targeting increasing the potential therapeutic efficacy of human CAR T cells as seen in our pre-clinical models.

**Figure 8.**
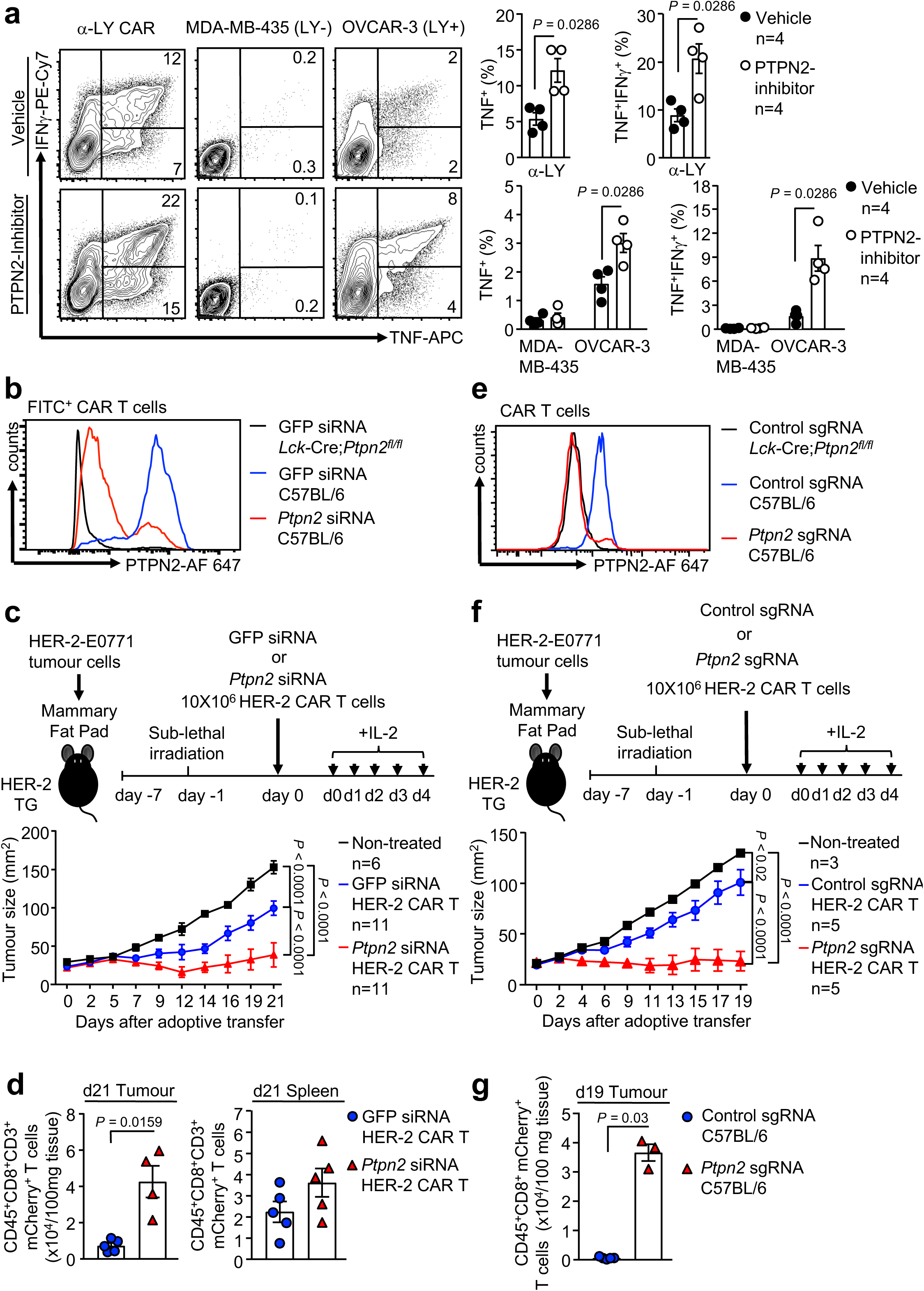
PTPN2 targeting enhances murine and human CAR T cell responses. **a**) CD8^+^ LY CAR T cells generated from human PBMCs were treated with PTPN2-inhibitor (+) or vehicle (-) followed by incubation with plate-bound α-LY, LY-negative MDA-MB-435 cells and LY-expressing OVCAR-3 cells. Cells were stained for CD8 and intracellular IFNγ and TNF, and the proportion of IFNγ^+^ and IFNγ^+^TNF^+^ CD8^+^CAR T cells determined by flow cytometry. **b**) HER-2 CAR T cells generated from C57BL/6 and *Lck*-Cre;*Ptpn2^fl/fl^* splenocytes were transfected with GFP versus *Ptpn2* siSTABLE™ FITC-conjugated siRNAs and intracellular PTPN2 levels were determined by flow cytometry. **c-d**) HER-2-E0771 mammary tumour cells (2×10^5^) were injected into the fourth inguinal mammary fat pads of female HER-2 TG mice. Seven days after tumour injection HER-2 TG mice received total body irradiation (4 Gy) followed by the adoptive transfer of 10×10^6^ HER-2 CAR T cells generated from C57BL/6 splenocytes transfected with GFP versus *Ptpn2* siSTABLE™ FITC-conjugated siRNAs two days before adoptive CAR T cell transfer. Mice were injected with IL-2 (50,000 IU/day) on days 0-4 after adoptive CAR T cell transfer and tumour growth was monitored. **d**) CD45^+^CD3^+^CD8^+^ mCherry^+^ CAR T cells numbers were determined in HER-2-E0771 positive tumours and spleens by flow cytometry 21 days post adoptive transfer. **e**) HER-2 CAR T cells generated from C57BL/6 and *Lck*-Cre;*Ptpn2^fl/fl^* splenocytes were transfected with Cas9 and control or *Ptpn2* sgRNAs using the Lonza 4D-Nucleofector and after two days intracellular PTPN2 levels determined by flow cytometry. **f-g**) HER-2-E0771 mammary tumour cells (2×10^5^) were injected into the fourth inguinal mammary fat pads of female HER-2 TG mice. Seven days after tumour injection HER-2 TG mice received total body irradiation (4 Gy) followed by the adoptive transfer of 10×10^6^ control HER-2 CAR T cells or those in which PTPN2 had been deleted by CRISPR RNP. Mice were injected with IL-2 (50,000 IU/day) on days 0-4 post adoptive CAR T cell transfer and tumour growth monitored. **g**) CD45^+^ CD8^+^ mCherry^+^ CAR T cells numbers were determined in HER-2-E0771 tumours by flow cytometry 19 days post adoptive transfer. Representative flow cytometry profiles and results (means ± SEM) from two independent experiments are shown. In (a, d, g) significance was determined using 2-tailed Mann-Whitney U Test. In (c, f) significance was determined using 2-way ANOVA Test.

### PTPN2 knockdown or deletion enhances the therapeutic efficacy of CAR T cells

Whole-body, T cell- or hematopoietic compartment-specific PTPN2 deletion in mice results in systemic inflammation, overt autoreactivity and morbidity ^5, 8, 12, 62, 63^. The potential for inflammatory complications, including cytokine release syndrome, preclude the utility of PTPN2 inhibitors for systemic therapy and the promotion of T cell mediated anti-tumour immunity. Accordingly, we sought alternate ways by which to target PTPN2 in the context of adoptive cell therapy. To this end we first knocked down *Ptpn2* in murine CAR T cells by RNA interference using nuclease resistant siRNA duplexes (siSTABLE™) that efficiently knock down genes for prolonged periods (Fig. 8b-d, Fig. S12c-f). HER-2 CAR T cells were transfected with GFP- or *Ptpn2*-specific siSTABLE™ siRNAs two days prior to adoptive cell therapy; PTPN2 was knocked down in approximately one third of total HER-2 CAR T cells as assessed by flow cytometry (**Fig. 8b**) using validated antibodies ^10^. *Ptpn2* knockdown enhanced the tumour antigen-specific activation/cytotoxic potential and killing capacity of HER-2 CAR T cells *ex vivo* (Fig. S12c-e) and markedly repressed the growth of HER-2-E0771 mammary tumours *in vivo* (**Fig. 8c**). The repression of tumour growth was accompanied by the significant infiltration of mCherry+ CD8+ HER-2 CAR T cells into tumours (**Fig. 8d**); infiltrating HER-2 T cells exhibited increased cytotoxic capacity (as assessed by IFNγ and TNF expression after PMA/ionomycin treatment *ex vivo*) (Fig. S12f). By contrast mCherry+ CD8+ HER-2 CAR T cell numbers in the spleen were not affected by *Ptpn2* knockdown (**Fig. 8d**). Therefore, the transient repression of PTPN2, even in a fraction of adoptively transferred CAR T cells, is sufficient to significantly enhance their activity/cytotoxicity and efficacy *in vivo*.

An alternate approach by which to target PTPN2 in T cells is through CRISPR-Cas9 genome-editing ^64^. In particular, we took advantage of Cas9 ribonucleoprotein (RNP)-mediated gene-editing to effectively delete PTPN2 in CAR T cells. This plasmid-free approach allows for efficient but transient genome editing *ex vivo* so that resultant CAR T cells do not overexpress Cas9 and do not elicit immunogenic responses post-adoptive transfer. To this end we transfected total CAR T cells with recombinant nuclear localized Cas9 pre-complexed with short guide (sg) RNAs capable of directing Cas9 to the *Ptpn2* locus (Fig. 8e-g; Fig. S12g). sgRNAs targeting the *Ptpn2* locus completely ablated PTPN2 protein in HER-2 CAR T cells (**Fig. 8e**) and enhanced their antigen-specific activation/cytotoxic potential (assessed by IFNγ production) and their capacity to specifically kill HER-2 expressing 24JK cells *ex vivo* (Fig. S12g).

Importantly we found that *Ptpn2* deletion increased the infiltration of mCherry+ HER-2 CAR T cells into HER-2-E0771 mammary tumours (**Fig. 8f**) and led to the effective eradication of HER-2-E0771 mammary tumours (**Fig. 8g**) *in vivo*. These results demonstrate that CRISPR-Cas9 genome editing can be used to efficiently ablate PTPN2 to enhance the therapeutic efficacy of CAR T cells in solid cancer.

## DISCUSSION

Approaches aimed at harnessing host immunity to destroy tumour cells have revolutionized cancer therapy ^1, 2^. Such approaches have relied on the targeting of immune checkpoints, such as PD-1, to alleviate inhibitory constraints on T cell-mediated anti-tumour immunity in immunogenic tumours, or alternatively on adoptive T cell therapy, especially that employing CAR T cells ^1, 2, 23^. Although the latter does not require pre-existing anti-tumour immunity, there are significant hurdles limiting efficacy and prohibiting the widespread utility of CAR T cells in the treatment of solid tumours ^23^. These include inefficient CAR T cell homing and infiltration into solid tumours and inadequate CAR T cell activation in the solid tumour microenvironment, which can be overtly immunosuppressive ^23^. Like PD-1 ^15, 17^, PTPN2 is fundamentally important in mediating T cell tolerance in mice and humans ^5, 10–14^. This is underscored by the striking phenotype similarities between PTPN2-deficient mice and those null for PD-1 ^5, 12, 15, 17, 62, 63^. Consistent with this, our studies herein demonstrate that the deletion of PTPN2 in T cells enhances cancer immunosurveillance and the anti-tumour activity of adoptively transferred T cells. In particular, our studies demonstrate that the deletion of PTPN2 not only drives the homing of CAR T cells to solid tumours, but also their activation to eradicate tumours in an otherwise immunosuppressive tumour microenvironment. Therefore, targeting PTPN2 may provide a means for enhancing the anti-tumour activity of T cells and extending the utility of CAR T cells beyond hematological malignancies to solid cancers.

A recent CRISPR loss-of-function screen in tumour cells identified PTPN2 as a top-hit for the recruitment of T cells and the sensitization of tumours to anti-PD-1 therapy ^55, 65^. This was reliant on PTPN2-deficiency driving the IFNγ-induced and STAT-1-mediated expression of antigen-presentation pathway genes and T cell chemoattractants, such as *Cxcl9* in tumour cells ^55^. In humans *CXCL9* expression is generally associated with increased CD8+ T cell infiltrates and improved overall survival and response to chemotherapy ^46^. Our own studies indicate that the deletion of PTPN2 in CAR T cells drives the expression of CXCR3 and the trafficking of CAR T cells to CXCL9/10-expressing mammary tumours. Indeed, we demonstrated that the homing and efficacy of PTPN2-deficient CAR T cells was reliant on tumours expressing STAT-1-driven CXCR3 chemokines. Thus, PTPN2-deficient CAR T cells might be especially effective against tumours such as estrogen receptor negative and triple negative breast cancers ^66^ or lung cancers ^67^ that have low PTPN2 levels. However it is important to note that a variety of human tumours, including for example breast, colon and ovarian cancers, can express CXCL9/10/11, or do so after chemotherapy ^46, 68–73^.

Although it may be possible to target PTPN2 systemically with drugs to enhance T cell and CAR T cell responses and potentially achieve synergistic effects through the targeting of PTPN2 in both tumour cells and T cells/CAR T cells, there are several important considerations. First, the currently available PTPN2 inhibitors target the active site of the enzyme and are therefore hydrophilic and have difficulties with bioavailability and pharmacokinetics ^74^. Second, we have shown that the inducible deletion of PTPN2 in the hematopoietic compartment of adult non-autoimmune-prone C57BL/6 mice is sufficient to promote the development of systemic inflammation and autoimmunity ^8^, whereas PTPN2 deletion in T cells in autoimmune-prone NOD1 mice markedly accelerates type 1 diabetes onset, as well as other autoimmune and inflammatory disorders, including colitis ^12^. Therefore, any systemic targeting of PTPN2 might exacerbate immune complications frequently associated with immunotherapy, including cytokine release syndrome that can be life-threatening in CAR T cell therapy ^23^. Third we and others have shown that the deletion of PTPN2 in some solid tumours can enhance tumorigenicity ^54, 66, 75^. For example, we have shown that PTPN2 deletion in the liver can facilitate the STAT-3-dependent development of hepatocellular carcinoma in obesity ^54^. Accordingly, we propose that the targeting of PTPN2 in adoptively transferred CAR T cells using stabilized siRNAs or CRISPR RNP genome-editing might be a safer and ultimately more effective means for enhancing the clinical efficacy of CAR T cells in solid tumours.

Our studies demonstrate that the deletion of PTPN2 enhances the function of CAR T cells by promoting antigen-induced LCK activation and cytokine-induced STAT-5 signalling. We and others have shown that LCK and STAT-5 can serve as *bona fide* substrates of PTPN2 in thymocytes/T cells ^5, 6, 9, 10, 38, 39^. In this study we found that correcting the increased LCK or STAT-5 signalling diminished the efficacy of PTPN2-deficient CAR T cells. Our studies indicate that the induction of LCK is necessary for the increased antigen-induced CAR T cell activation, IL-2/IL-15 receptor subunit (CD25, CD122 and CD132) expression and cytotoxicity of PTPN2-deficent CAR T cells, whereas the concomitant direct promotion of STAT5 phosphorylation and cytokine signalling might not only facilitate the acquisition of CTL activity, but also promote homing to CXCL9/10/11 expressing tumours through the induction of CXCR3. IL-2-induced STAT-5 signalling is necessary for the differentiation of activated CD8+ T cells into effectors and the acquisition of CTL activity ^49, 50^. Although the precise mechanism by which STAT-5 promotes CXCR3 in PTPN2-deficient CAR T cells remains unclear, CXCR3 is elevated in effector CD8+ T cells and previous studies have shown that STAT-5 can drive T-bet and Eomes expression ^51^. Consistent with this our studies demonstrate that the elevated STAT-5 signalling in PTPN2-deficient CAR T cells was also associated with increased T-bet expression. Importantly, T-bet can drive the expression of CXCR3 ^52^ and both T-bet and Eomes have been shown to be important for the tumour infiltration and anti-tumour activity of cytotoxic CD8+ T cells ^53^. Moreover, in the context of viral infection, CXCR3 on CD8+ T cells facilitates T cell homing to infected tissue and the ability of T cells to locate an eliminate infected cells ^76^. In our studies we found STAT-5 and the resultant increased CXCR3 promoted the homing of PTPN2-deficient CAR T cells to CXCL9/10-expressing mammary tumours within 3 days of adoptive transfer. By contrast significant PTPN2-deficient CAR T cells were not detected in the HER-2+ cerebellum that did not express CXCL9/10. As systemic inflammation, tissue damage, autoimmunity and morbidity were not evident in HER-2 transgenic mice, despite the abundant HER-2 expression in the cerebellum, we conclude that the enhancement of STAT5 signalling limits ‘on-target off-tumour’ toxicities by promoting the specific homing of CAR T cells to CXCL9/10 expressing tumours.

The results of this study have established the importance of PTPN2 in T cell immunosurveillance and defined a novel target for bolstering the anti-tumour activity of T cells, especially in the context of adoptive T cell therapy. In particular, our findings have defined a novel approach by which to enhance the efficacy of CAR T cells and extend their utility to the treatment of solid tumours without promoting systemic inflammation/autoimmunity and morbidity.

## ACKNOWLEDGMENTS

This work was supported by the National Health and Medical Research Council (NHMRC) of Australia (to T.T., T.G., P.K.D.), Cancer Council Victoria (F.W.) and the NIH (RO1 CA207288 to Z.Y.); T.T. and P.K.D are NHMRC Research Fellows, K.H. a Rhian and Paul Brazis Fellow in Translational Melanoma Immunology, T.G. a Sylvia & Charles Viertel Charitable Foundation Senior Biomedical Research Fellow and J.W. a Cancer Council WA Fellowship. S.L.P. is supported by an Elizabeth and Vernon Puzey Postgraduate Scholarship (University of Melbourne).

## ONLINE METHODS

### Cell lines and mice

The C57BL/6 mouse breast carcinoma cell line E0771 (a gift from Robin Anderson, Peter MacCallum Cancer Centre) ^77^ and the C57BL/6 mouse sarcoma cell line 24JK (a gift from Patrick Hwu, NIH, Bethesda, Maryland, USA) ^78^ were genetically engineered to express truncated human HER-2 (HER-2-E0771) as described previously ^79^. The C57BL/6 mouse mammary tumour cell line AT-3 was genetically engineered to express chicken ovalbumin (AT-3-OVA) and has been previously described ^80^. The GP+E86 packaging line was generated as previously described ^81^. The human epithelial adenocarcinoma cell line OVCAR-3 (ATCC^®^HTB-161^™^) and the human melanoma cell line MDA-MB-435 (ATCC^®^HTB-219^™^) were obtained from the ATCC, Manassas, Virginia, USA. HER-2-E0771 cells were engineered to inducibly overexpress murine PTPN2 (HER-2-E0771-PTPN2^hi^) in response to doxycycline using the Tet-On 3G Inducible Expression System according to the manufacturer’s instructions (Clontech). Tumour cells were cultured in RPMI 1640 (HER-2-E0771, OVCAR-3, MDA-MB-435) or high-glucose DMEM (AT3-OVA) supplemented with 10% FBS, L-glutamine (2mM), penicillin (100 units/mL)/streptomycin (100 μg/mL), MEM non-essential amino acids (0.1 mM), sodium-pyruvate (1 mM), HEPES (10 mM) and 2-mercaptoethanol (50 μM).

Mice were maintained on a 12 h light-dark cycle in a temperature-controlled high barrier facility with free access to food and water. 6-10 week old female Ly5.1 (B6.SJL-*Ptprc^a^Pepc^b^*/BoyJ) and human HER-2 transgenic (TG) recipient mice and 6-8 week old female donor mice were used for adoptive transfers. For *ex-vivo* experiments either male or female mice were used. Aged- and sex-matched littermates were used in all experiments. *Ptpn2^fl/fl^* and *Lck*-Cre;*Ptpn2^fl/^*^fl^ mice and the corresponding OT-1 TCR transgenic mice were described previously ^5^. *p53^+/–^* (C57BL/6) mice have been described previously ^26^ and were a gift from Prof. Andreas Strasser (WEHI, Melbourne, Australia). *Ptpn2^fl/fl^*;*p53^+/–^* and *Lck*-Cre;*Ptpn2^fl/fl^;p53^+/–^* mice were generated by crossing *p53^+/–^* (C57BL/6) with *Lck*-Cre;*Ptpn2^fl/fl^* mice. B6.129S2-*Lck*/J mice were a gift from Dr Andre Veillette (McGill University, Montreal) and were bred with *Lck*-Cre;*Ptpn2^fl/fl^* mice to generate *Lck*-Cre;*Ptpn2^fl/fl^*;*Lck^+/–^* mice. Ly5.1 mice and C57BL/6 were purchased from the WEHI Animal Facility (Kew, Australia) and human HER-2 (C57BL/6) transgenic (TG) mice were bred and maintained at the Peter MacCallum Cancer Centre.

## Materials

For cell stimulation, anti-CD3ε (clone 145-2C11) and anti-CD28 (clone 37.51) antibodies were purchased from BD Biosciences. Antibodies against p-(Y701) STAT1 (clone 58D6) and STAT1 were from Cell Signaling. Anti-Actin (clone ACTN05) was purchased from Thermo Fisher Scientific. The mouse antibody against PTPN2 (clone 6F3) was provided by M. Tremblay (McGill University). For immunofluorescence staining and immunohistochemistry rabbit anit-CD3ε and mouse anti-HER-2 from Abcam was used. Recombinant human IL-2, murine IL-7 and IL-15 used for T cell stimulation or IFNγ used for stimulating tumour cells were purchased from the NIH or PeproTech, respectively. Retronectin was purchased from Takara, Dnase I and doxycycline from Sigma-Aldrich, the mouse anti-nuclear antibodies Ig’s (total IgA+G+M) ELISA Kit from Alpha Diagnostic Int., the Transaminase II Kit from Wako Pure Chemicals, FBS from Thermo Scientific, Dulbecco-Phosphate Buffered Saline (D-PBS), RPMI 1640, DMEM, MEM non-essential amino acids and sodium-pyruvate from Invitrogen and collagenase type IV from Worthington Biochemical.

### Flow cytometry

Single cell suspensions from spleen and lymph nodes were obtained by mashing excised organs through 40 μM cell strainers (BD Biosciences). Hepatic lymphocytes were isolated from liver tissue and mashed through a wire mesh (200 μM pore size). Hepatocytes were removed by Percoll 33.75% (v/v) (GE Healthcare) gradient centrifugation at room temperature. Cells were incubated for 7 minutes on ice in Red Blood Cell Lysing Buffer Hybri-Max™ (Sigma-Aldrich) for the removal of contaminating erythrocytes. Lymphocyte counts (4-15 μm) were determined with a Z2 Coulter Counter (Beckman Coulter). For surface staining, cells (1×10^6^/10 µl) were resuspended in D-PBS supplemented with 2% (v/v) FBS containing the antibody cocktail in 96-well microtiter plates (BD Biosciences) for 20 min on ice in the dark. Cells were fixed and permeabilized with the BD Cytofix/Cytoperm kit according to the manufacturer’s instructions. For the detection of intracellular FoxP3 the Foxp3/Transcription Factor Staining Buffer Set (eBioscience) was used according to the manufacturer’s instructions. For the detection of intracellular p(Y418)-SFK and PTPN2, cells were fixed in 200 μl Cytofix™ Fixation Buffer (BD Biosciences) for 15 min at 37°C. Cells were washed twice with D-PBS to remove excess paraformaldehyde and permeabilized in 200 μl methanol (90% v/v) for p(Y418)-SFK detection or in 200 μl methanol/acetone (50:50) for PTPN2 detection at -20°C and processed for flow cytometry on the following day. Secondary antibodies against mouse IgG (H+L) F(ab’)2 fragment conjugated to AlexaFluor 647 or FITC (Molecular Probes, Thermo Fisher Scientific) was used for the detection of PTPN2. Cells were stained with the specified intracellular antibodies on ice for 30 min and analyzed using a LSRII, Fortessa, Symphony (BD Biosciences) or CyAn™ ADP (Beckman Coulter).

For the detection of serum cytokines either the LEGENDplex T_h_ Cytokine Panel™ from BioLegend or the BD CBA Mouse Inflammation Kit™ were used according to the manufacturers’ instructions.

For FACS sorting, cells were stained in 15 mL Falcon tubes (BD Biosciences) for 30 min on ice and purified using either a BD Influx cell sorter, or the BD FACSAria II, BD FACSAria Fusion 3 or BD FACSAria Fusion 5 instruments.

Data was analyzed using FlowJo8.7 or FlowJo10 (Tree Star Inc.) software. For cell quantification, a known number of Calibrite™ Beads (BD Biosciences) or Nile Red Beads (Prositech) or Flow-Count Fluorospheres (Beckman Coulter) were added to samples before analysis.

The following antibodies from BD Biosciences, BioLegend or eBioscience were used for flow cytometry: Phycoerythrin (PE) or peridinin-chlorophyll cyanine 5.5 (PerCP-Cy5.5)-conjugated CD3 (145-2C11); PerCP-Cy5.5 or phycoerythrin-cyanine 7 (PE-Cy7)-conjugated CD4 (RM4-5); BV711, Pacific Blue-conjugated (PB), allophycocyanin (APC)-Cy7 or APC-conjugated CD8 (53-6.7); PerCP-Cy5.5, PE or APC-Cy7-conjugated CD25 (P61); Fluorescein isothiocyanate (FITC), V450, AlexaFluor 700 or PE-Cy7-conjugated CD44 (IM7); APC-conjugated CD45 (30-F11); V450, APC, AlexaFluor 488 or PE-conjugated CD45.1 (Ly5.1; A20); FITC, PB or PerCP-Cy5.5-conjugated CD45.2 (Ly5.2; 104); APC-conjugated CD45R (B220; RA3-6B2); PE-Cy7, APC or PE-conjugated CD62L (Mel14); PE-conjugated CD122 (TM-β1); PE-conjugated CD132 (4G3); AlexaFluor 647-conjugated CD183 (CXCR3-173); PerCP-eFluo710-conjugated CD185 (CXCR5; SPRCL5); PE-Cy7 conjugated CD197 (CCR7; 4B12); PE-conjugated CD223 (LAG-3; C9B7W); PE-Cy7-conjugated CD279 (PD-1, RMP1-14); FITC-conjugated GL-7 (GL-7); PE-Cy7 conjugated CD49d (9C10); APC-conjugated KLRG-1 (2F1); FITC-conjugated CD11b (M1/70); PE-conjugated Ly-6G/Ly-6C (Gr-1; RB6-8C5); APC-Cy7-conjugated Ly6C (AL-21); PE-conjugated Ly-6G (clone 1A8), APC-conjugated F4/80 (BM8); BV421-conjugated CD206 (C068C2); PE-Cy7-conjugated IFNγ (XMG1.2); FITC or APC-conjugated TNF (MP6-XT22); AlexaFluor 647-conjugated Granzyme B (GB11), PE-Cy7-conjugated TCR-Vα2 (B20.1); eFluor 660-conjugated p(Y418)-Src (SC1T2M3); AlexaFluor 647-conjugated T-bet (4B10), (V450-conjugated FoxP3 (clone MF23).

The following antibodies from Miltenyi Biotec were used for flow cytometry: PE-conjugated CD197 (CCR7; REA108); VioBright FITC-conjugated CD8 (BW135/80); VioBlue-conjugated CD45RA (REA1047).

### Generation of murine CAR T cell

Splenocytes were isolated from murine spleens by mechanical disruption and contaminating red blood cells were removed by incubation with 1 ml of Blood Cell Lysing Buffer Hybri-Max™ (Sigma-Aldrich) for 7 min at room temperature per spleen. Lymphocytes (0.5×10^7^/ml) were cultured overnight with anti-CD3ε (0.5 µg/ml) and anti-CD28 (0.5 µg/ml) in the presence of 100 IU/ml human IL-2 and 2 ng/ml murine IL-7 in complete T cell medium [RPMI supplemented with 10% FBS, L-glutamine (2mM), penicillin (100 units/mL)/streptomycin (100 μg/mL), non-essential amino acids, Na-pyruvate (1 mM), HEPES (10 mM) and 2-mercaptoethanol (50 μM)]. Dead cells were removed by Ficoll centrifugation (GE Healthcare) according to the manufacturer’s instructions. Retrovirus encoding a second generation chimeric antibody receptor (CAR) consisting of an extracellular scFv–anti–human HER-2, a membrane proximal CD8 hinge region and the transmembrane and the cytoplasmic signaling domains of CD28 fused to the cytoplasmic region of CD3ζ (scFv-anti-HER-2-CD28-ζ) was obtained from the supernatant of the GP+E86 packaging line as described previously ^35^. Retroviral supernatant was added to non-treated retronectin-coated (10 µg/ml) 6-well plates (Takara Bio) and spun for 30 min (1,200 x g) at room temperature. T cells were resuspended in 1 ml of additional retroviral-containing supernatant supplemented with IL-2 and IL-7 and then added to the retronectin-coated plates to a final volume of 5 ml/well with a final T cell concentration of 1 × 10^7^ per well. T cells were spun for 90 min (1,200 x g) at room temperature and incubated overnight before the second viral transduction. T cells were maintained in IL-2– and IL-7–containing media and cells used at days 7–8 after transduction.

### Generation of human CAR T cells

Human peripheral blood mononuclear cells (PBMCs) were isolated from normal donor buffy coats. PBMCs were stimulated with anti-human CD3 (OKT3; 30 ng/ml, Ortho Biotech) and human IL-2 (600 IU/ml) in complete T cell medium. Retrovirus encoding a second generation chimeric antibody receptor (CAR) consisting of an extracellular scFv-anti-human LeY domain, a membrane proximal CD8 hinge region and the transmembrane and the cytoplasmic signaling domains of CD28 fused to the cytoplasmic region of CD3ζ was obtained from the supernatant of the PG13 packaging cell line and has been described previously ^60, 61^. Retroviral supernatant (5 ml) was added to non-treated retronectin-coated (10 µg/ml) 6-well plates (Takara Bio) and incubated for 4 h at 37°C. Supernatant was removed and 2.5×10^6^ T cells were incubated overnight in 5 ml of complete T cell medium supplemented with human IL-2 before the second viral transduction. At day 7 post retroviral transduction, human CAR T cells were sorted for CD34^+^ (CD34 MicroBead Kit UltraPure, MACS, Miltenyi Biotec) CAR T cells and expanded for up to 15 days in the presence of human IL-2. Expression of the chimeric receptor was determined by staining with AlexaFlour 647-conjugated anti-3S193 idiotype [supplied by Ludwig Institute for Cancer Research (LICR)] and truncated CD34 was detected with PE-conjugated anti-human CD34 (BioLegend). Cell surface phenotyping of transduced cells was determined by staining with BV785-conjugated anti-human CD3 (UCHT1, BioLegend), BV605-conjugated anti-human CD4 (OKT-4, BioLegend) and PE-Cy7-conjugated anti-human CD8 (SK1, BioLegend).

### Assessment of cytokine production in human CAR T cells

CAR T cells (2×10^5^) generated from human PBMC were incubated with Lewis Y expressing OVCAR-3 cells (1×10^5^) or Lewis Y negative MDA-MB-435 cells (1×10^5^) or plate-bound anti-LY (1 μg/ml; Cell Therapies Pty Ltd, Peter MacCallum Cancer Centre) for 6 h at 37°C in complete T cell medium. GolgiPlug™ and GolgiStop™ were added 3 h before cells were processed for surface and intracellular staining for CD4 (BV785-conjugated anti-human; OKT-4, BioLegend), CD8 (APC-H7-conjugated anti-human; SK1, BD Bioscience), IFNγ (PE-Cy7-conjugated anti-human; B27, BioLegend) and TNF (APC-conjugated anti-human TNF; MAb11; BioLegend) by flow cytometry.

### Cytokine signalling in CAR T cells

CAR T cells generated from mouse splenocytes were incubated with plate-bound anti-CD3ε for 24 h at 37°C in complete T cell medium. Cells were washed twice and rested in RPMI/1%FBS for 1 h. Cells were stimulated for the indicated time points with mouse recombinant IL-2 (5 or 10 ng/ml) or IL-15 (5 or 10 ng/ml) in 100 μl RPMI/1%FBS. Cell were fixed in 100 μl Cytofix™ Fixation Buffer (BD Biosciences) for 15 min at 37°C. Cells were washed twice with D-PBS to remove excess paraformaldehyde and permeabilized in 200 μl methanol/acetone (50:50) overnight at -20°C. Permeabilized cells were washed three times with D-PBS/5% FBS and processed for intracellular staining for p(Y694)-STAT-5 (D47E 7 XP^®^ rabbit, Cell Signaling Technology) in D-PBS/5% FBS for 1 h at room temperature. Secondary antibodies against rabbit IgG (H+L) F(ab’)2 fragment coupled to DyLight 649 (Jackson ImmunoResearch) were used for staining for p(Y694)-STAT-5 detection.

### CAR T cell proliferation

CAR T cells generated from mouse splenocytes were stained with fluorochrome-conjugated antibodies for CD8, CD62L and CD44 and sorted for CD8^+^CD44^hi^CD62L^hi^ CAR T cells. For the assessment of CAR T cell proliferation by CellTrace™ Violet (CTV; Molecular Probes, Thermo Fisher Scientific) dilution, purified CD8^+^CD44^hi^CD62L^hi^ CAR T cells were incubated with CTV in D-PBS supplemented with 0.1% (v/v) BSA at a final concentration of 2 μM for 10 min at 37°C. Cells were then washed three times with D-PBS supplemented with 10% (v/v) FBS. CAR T cells (2×10^5^) were incubated with HER-2 expressing or HER-2 negative 24JK sarcoma cells (1×10^5^) in complete T cell medium.

For the assessment of CAR T cell proliferation after anti-CD3ε stimulation, purified CD8^+^CD44^hi^CD62L^hi^ CAR T cells were incubated with plate-bound anti-CD3ε for 24 h at 37°C in complete T cell medium. Pre-activated CAR T Cells (2×10^5^) were labelled with 5 μM CTV and incubated with HER-2 expressing or HER-2 negative 24JK sarcoma cells (1×10^5^) in complete T cell medium. At various time points, proliferating cells were harvested and CTV dilution was monitored by flow cytometry.

For cell quantification, Nile Red Beads (Prositech) or Flow-Count Fluorospheres (Beckman Coulter) were added to samples before analysis.

### PTPN2-inhibition studies

T cells were incubated rocking on an orbital shaker with PTPN2-inhibitor Compound 8 (100 nM) or vehicle (DMSO 1% v/v) in serum-free RPMI 1640 for 1 h at 37°C. Cells were washed three times with complete T cell medium and processed for T cell activation studies.

### Treatment of HER-2-E0771 tumour-bearing HER-2 TG mice

Female human HER-2 TG mice were anaesthetized with Ketamine (100 mg/kg) and Xylazil (10 mg/kg) (Troy Laboratories) and injected orthotopically with 2 × 10^5^ HER-2-E0771 cells resuspended in 20 µl D-PBS into the fourth mammary fat pad. At day 6 post tumour cell injection, human HER-2 TG mice were preconditioned with total-body irradiation (4 Gy) prior to the adoptive transfer of 6 × 10^6^ FACS-purified CD8^+^CD44^hi^CD62L^hi^ CAR T cells. Mice were treated with 50,000 IU IL-2 on days 0– 4 after T cell transfer.

### Treatment of AT-3-OVA tumour bearing Ly5.1 mice

Female Ly5.1 mice were anaesthetized with Ketamine (100 mg/kg) and Xylazil (10 mg/kg) (Troy Laboratories) and were injected orthotopically with 1×10^6^ AT-3-OVA cells resuspended in 20 µl D-PBS into the fourth mammary fat pad. At day 7 post tumour injection, 2 × 10^6^ FACS-purified CD8^+^CD44^hi^CD62L^hi^ OT-1 T cells were adoptively transferred.

### Assessment of autoimmunity in HER-2 TG mice

Serum liver enzymes aspartate aminotransferase (AST/GOT) and alanine aminotransferase (ALT/GPT) were determined with the Transaminase CII kit from Wako Pure Chemical. Serum (20 μl) from HER-2 TG mice or GOT/GPT standards (20 1. μl) were incubated with GOT or GPT enzyme (500 μl) at 37°C for 5 min. The colour forming solution (500 μl) was added and the reaction terminated with stopping solution (2 ml) after 20 min. Absorbance was read at 555 nM and Karmen units/ml were calculated as sample (OD555 nm)/standard (OD555 nm) * dilution factor * 100 U/ml.

Serum anti-nuclear antibodies were detected with the mouse anti-nuclear antibodies Ig’s (total IgA+G+M) ELISA Kit from Alpha Diagnostic International. Serum (1:20 dilution) was incubated on 8-well polystyrene strips coated with ENA (extractable nuclear antigens) at room temperature for 1 h. Plates were washed four times and incubated with Streptavidin-HRP for 30 min followed by five washes. 3,3’,5,5’-tetramethylbenzidine was added and reactions developed for 15 min at room temperature in the dark and stopped with 1% sulphuric acid. Absorbance was read at 450 nM and activity units were calculated as sample (OD450 nm)/standard (OD450 nm) * dilution factor * 200 U/ml.

### Analysis of tumour-infiltrating T cells

Tumour bearing mice were sacrificed and tumours were excised and digested at 37°C for 30 min using a cocktail of 1 mg/ml collagenase type IV (Worthington Biochemicals) and 0.02 mg/ml DNase (Sigma-Aldrich) in DMEM supplemented with 2% (v/v) FBS. Cells were passed through a 70 µm cell strainer (BD Biosciences) twice and processed for flow cytometry. For the detection of intracellular cytokines in tumour infiltrating lymphocytes, cells (1×10^6^/200 µl) were stimulated with PMA (20 ng/ml; Sigma-Aldrich) and ionomycin (1 µg/ml; Sigma-Aldrich) in the presence of GolgiPlug™ and GolgiStop™ (BD Biosciences) for 4 hours at 37°C in complete T cell medium.

### Re-challenge of tumour-infiltrating T cells

AT-3-OVA tumour bearing C57BL/6 mice were sacrificed and tumours were collected in ice cold D-PBS and then digested with 50 µg/ml Liberase (Roche Diagnostics, Indianapolis, IN) in Hank’s Balanced Salt Solution (Ca^2+^, Mg^2+^) and 10 mM HEPES for 1h at 37°C. Cells were passed through a 70 µm cell strainer twice (BD Biosciences, San Jose, CA) and washed three times with 50 ml PBS. Cells were resuspended in DMEM supplemented with 10% FBS, 2mM L-glutamine, penicillin (100 units/ml), streptomycin (100 μg/ml) and seeded into 24-well plates. Cells were incubated at 37°C until they reached 80% confluency. Tumour-infiltrating T cells from AT-3-OVA tumour bearing C57BL/6 mice *Ptpn2^fl/fl^* and *Lck*-Cre;*Ptpn2^fl/fl^* mice were labeled with 2 μM CTV and incubated with adherent, 80% confluent AT-3-OVA tumour cells overnight. GolgiPlug™ and GolgiStop™ were added 3 h before cells were processed for intracellular staining for IFNγ and TNF by flow cytometry.

### Cytotoxic T cell assays

For the assessment of CAR T cell cytotoxicity, HER-2 expressing 24JK sarcoma cells (5 mM CTV) and HER-2 negative 24JK sarcoma cells (0.5 mM CTV) were labelled with CellTrace™ Violet (CTV; Invitrogen-Molecular Probes) in D-PBS supplemented with 0.1% (v/v) BSA for 15 min at 37°C. Tumour cells were then washed three times with D-PBS supplemented with 10% (v/v) FBS and mixed at a 1:1 ratio. FACS-purified CD8^+^ central memory (CD44^hi^CD62L^hi^) or effector/memory (CD44^hi^CD62L^lo^) or total HER-2-specific CAR T cells were added at different concentrations to the mix of HER-2-expressing (5×10^4^) and HER-2 negative (5×10^4^) 24JK sarcoma cells and incubated for four hours at 37°C in complete T cells medium. Antigen-specific target cell lysis (24JK-HER-2 cell depletion) was monitored by flow cytometry.

### Immunohistochemistry

For brain immunohistochemistry, mice were perfused transcardially with heparinized saline [10,000 units/l heparin in 0.9% (w/v) NaCl] followed by 4% (w/v) paraformaldehyde in phosphate buffer (0.1 M, pH 7.4). Brains were post-fixed overnight and then kept for four days in 30% (w/v) sucrose in 0.1 M phosphate buffer to cryoprotect the tissue, before freezing on dry ice. 30 μm sections (120 μm apart) were cut in the coronal plane throughout the entire rostral-caudal extent of the cerebellum. For detection of CD3ε, sections were subjected to antigen retrieval in citrate acid buffer [10 mM Sodium Citrate, 0.05% (v/v) Tween 20, pH 6.0] at 85°C for 20 min. Sections were incubated at room temperature for 2 h in blocking buffer [0.1M phosphate buffer, 0.2% (v/v) Triton X-100, 10% (v/v) normal goat serum (Sigma-Aldrich) and then overnight at 4°C in rabbit anti-CD3 (Dako) in 1% (v/v) blocking buffer. After washing with PBS, sections were incubated with goat anti-rabbit Alexa Fluor 488 conjugated secondary antibody (Life Technologies) in blocking buffer for 2 h at room temperature. Sections were mounted with Mowiol 4-88 mounting media and visualized using an Olympus Provis AX70 microscope. Images were captured with an Olympus DP70 digital camera and processed using AnalySIS (Olympus) software. To assess gross tissue morphology, a subset of sections throughout the entire rostral-caudal extent of the cerebellum were stained with hematoxylin and eosin.

For detection of HER-2, 30 μm sections throughout the entire rostral-caudal extent of the cerebellum from C57BL/6 or HER-2 TG C57BL/6 mice were subjected to antigen retrieval and incubated in blocking buffer (as described above). Sections were then incubated overnight in (4°C) with anti-c-ErbB2/c-Neu (Ab-3) mouse mAb (3B5) (1:500, Calbiochem). After washing with PBS, HER-2-positive cells were visualized using rabbit IgG VECTORSTAIN ABC Elite and DAB (3,3’-diaminobenzidine) Peroxidase Substrate Kits (Vector Laboratories, UK) and visualized using a bright field microscope.

For mammary fat pad tumour immunohistochemistry animals were culled and mammary fat pad immediately dissected and fixed in buffered formalin solution for 48 h. Tissues were embedded in paraffin and 4 µm sections of the entire block prepared. Every tenth to fourteenth section of the tissue was used to detect HER-2 and CD3 by immunohistochemistry. After deparaffinization and rehydration sections were subjected to antigen retrieval in Tris/EDTA buffer (pH8.0) at 120 °C for 10 min. Sections were blocked with 5% (v/v) bovine serum albumin in 0.1 M phosphate buffer for 1 h at room temperature and incubated overnight (4°C) with anti-c-ErbB2/c-Neu (Ab-3) mouse mAb (3B5) (1:500, Calbiochem) or anti-CD3 (1:500; RAM 34 clone, 14-0341, Affymetrix eBioscience, San Diego, CA). After washing with PBS, HER-2 and CD3-positive cells were visualized using rabbit IgG VECTORSTAIN ABC Elite and DAB (3,3’-diaminobenzidine) Peroxidase Substrate Kits (Vector Laboratories, UK) and counterstained with hematoxylin. Sections were visualized on a Zeiss Axioskop 2 mot plus microscope (Carl Zeiss, Göttingen, Germany).

### Quantitative real-time PCR

RNA was extracted with TRIzol reagent (Thermo Fisher Scientific, #15596018) and RNA quality and quantity determined using a NanoDrop 2000 (Thermo Fisher Scientific). mRNA was reverse transcribed using a High-Capacity cDNA Reverse Transcription Kit (Applied Biosystems, # 4368814) and processed for quantitative real-time PCR either using the Fast SYBR™ Green Master Mix (Applied Biosystems, # 4385612). Primer sets from PrimePCR™ SYBR® Green Assay (Bio-Rad, #10025636) were utilized to perform quantitative PCR detecting *Cxcl9, Cxcl10, Tgfb1*, *Cd274*, *H2-K1*, and *Rps18*. Primer sets for *HER-2, Nono, and SerpinB14 (ovalbumin)* were purchased from Sigma-Aldrich. Relative gene expression (ΔCt) was determined by normalisation to the house-keeping gene *Nono* throughout the study except for Figure 2C, where *Rps18* was used and ΔΔCt analysis performed.

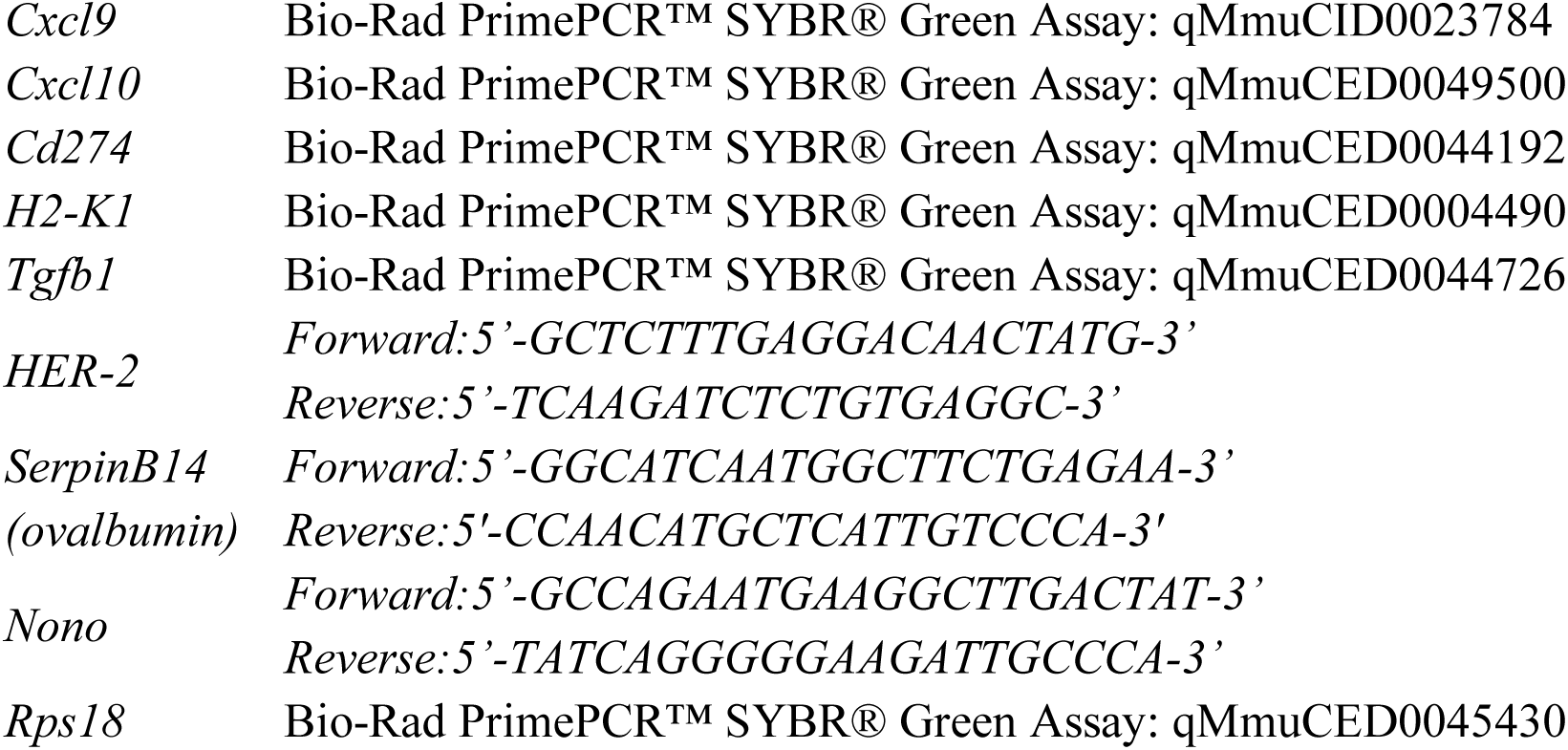

### Ptpn2 knock down using siSTABLE™ siRNAs targeting in murine CAR T cells

*Ptpn2* was knocked down transiently in HER-2 CAR T cells using nuclease resistant *Ptpn2* siRNA duplexes (*Ptpn2*; GCCCAUAUGAUCACAGUCG) (siSTABLE™; Dharmacon Thermo Scientific); siRNA duplexes (siSTABLE™) for enhanced green fluorescent protein (GFP; CAAGCUGACCCUGAAGUUC) were used as a control. CAR T cells were transfected with *Ptpn2* siRNA (300 nM) conjugated to FITC or GFP siRNA (300 nM) conjugated to FITC two days prior to adoptive T cell therapy using the Mouse T cell Nucleofector™ Kit (Lonza Bioscience) according to the manufacturer’s instructions.

### CRISPR-Cas9 genome-editing in murine CAR T cells

*Ptpn2* was deleted in HER-2 CAR T cells using Cas9 ribonucleoprotein (RNP)- mediated gene-editing. Briefly, total CAR T cells were transfected with recombinant Cas9 (74 pmol; Alt-R S.p. Cas9 Nuclease V3, IDT) pre-complexed with short guide (sg) RNAs (600 pmol; Synthego) targeting the *Ptpn2* locus (*Ptpn2*; 5’-AAGAAGUUACAUCUUAACAC) or non-targeting sgRNAs (GCACUACCAG AGCUAACUCA) as a control two days prior to adoptive T cell therapy using the P3 Primary Cell 4D-Nucleofector X™ Kit (Lonza Bioscience) according to the manufacturer’s instructions.

### Overexpression of Ptpn2 in HER-2-E0771 cells

The Tet-on 3G inducible expression system (Clonetech Laboratories) was used to generate the HER-2-E0771-PTPN2^hi^ cell line. Briefly, murine *Ptpn2* cDNA from HER-2-E0771 cells was reverse transcribed and amplified by PCR and cloned into the *Sma*I and *Eco*R1 restriction sites of Tre-3G plasmid (Clontech Laboratories) to generate the Tre-3G-*Ptpn2* construct; the fidelity of the cloned cDNA was confirmed by dideoxy sequencing. HEK293T cells were transfected with either the Tre-3G-*Ptpn2 or* Tet-On 3G constructs using the Lenti-X Packing system (Clontech Laboratories) according to the manufacturer’s instructions. HER-2-E0771 mammary tumour cells were transduced first with Tet-On 3G lentivirus and selected with G418 (0.8 mg/ml) and subsequently transduced with Tre-3G-*Ptpn2* lentivirus and selected with puromycin (1.5 μg/ml). Where indicated the resultant HER-2-E0771-PTPN2^hi^ cells were incubated with 2 mg/ml doxycycline (DOX) to induce the expression of PTPN2. Cells were serum starved in DMEM medium without FBS for 12 h and stimulated with 2 ng/ml IFNγ for the indicated times and processed for immunoblotting or incubated with 1 ng/ml IFNg for 24 h and processed for quantitative real time PCR.

For *in vivo* studies 2 × 10^5^ HER-2-E0771-PTPN2^hi^ cells were resuspended in 20 µl D-PBS and injected orthotopically into the fourth mammary fat pad of female human HER-2 TG mice. At day 5 post tumour cell injection mice were administered DOX (2 mg/ml) in the drinking water for the entirety of the experiment. At day 6 post tumour cell injection, human HER-2 TG mice were pre-conditioned with total-body irradiation (4 Gy) prior to the adoptive transfer of 6 × 10^6^ FACS-purified CD8^+^CD44^hi^CD62L^hi^ CAR T cells. Mice were treated with 50,000 IU IL-2 on days 0–4 after T cell transfer.

### Statistical analyses

Statistical analyses were performed with Graphpad Prism software 7.0b using the non-parametric using 2-tailed Mann-Whitney U Test, the parametric 2-tailed Student’s t test, the 1-way or 2-way ANOVA-test using Turkey or Sidak post-hoc comparison or the Log-rank (Mantel-Cox test) where indicated. p<0.05 was considered as significant.

### Animal ethics

All experiments were performed in accordance with the NHMRC Australian Code of Practice for the Care and Use of Animals. All protocols were approved by the Monash University School of Biomedical Sciences Animal Ethics Committee (Ethics number: MARP/2012/124) or the Peter MacCallum Animal Ethics and Experimentation Committee (Ethics numbers: E570, E582 and E604).

---

**Supplementary figure 1.**
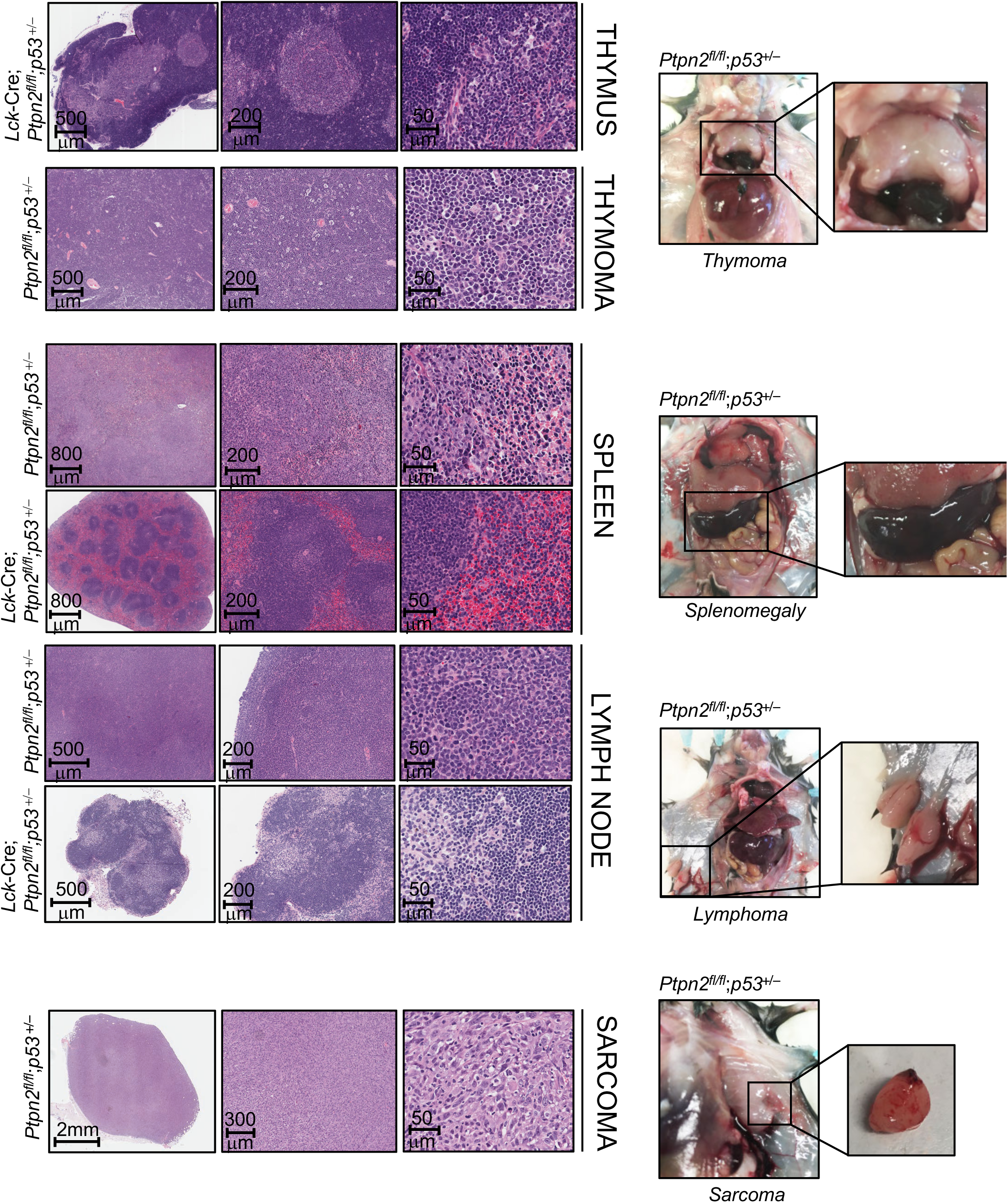
Tissue architecture and tumours in p53^+/–^ mice. Gross images and histological analyses (Hematoxylin and Eosin: H&E) from 12 month old *Ptpn2^fl/fl^*;*p53^+/–^* and *Lck*-Cre;*Ptpn2^fl/f^;p53^+/–^* mice.

**Supplementary figure 2.**
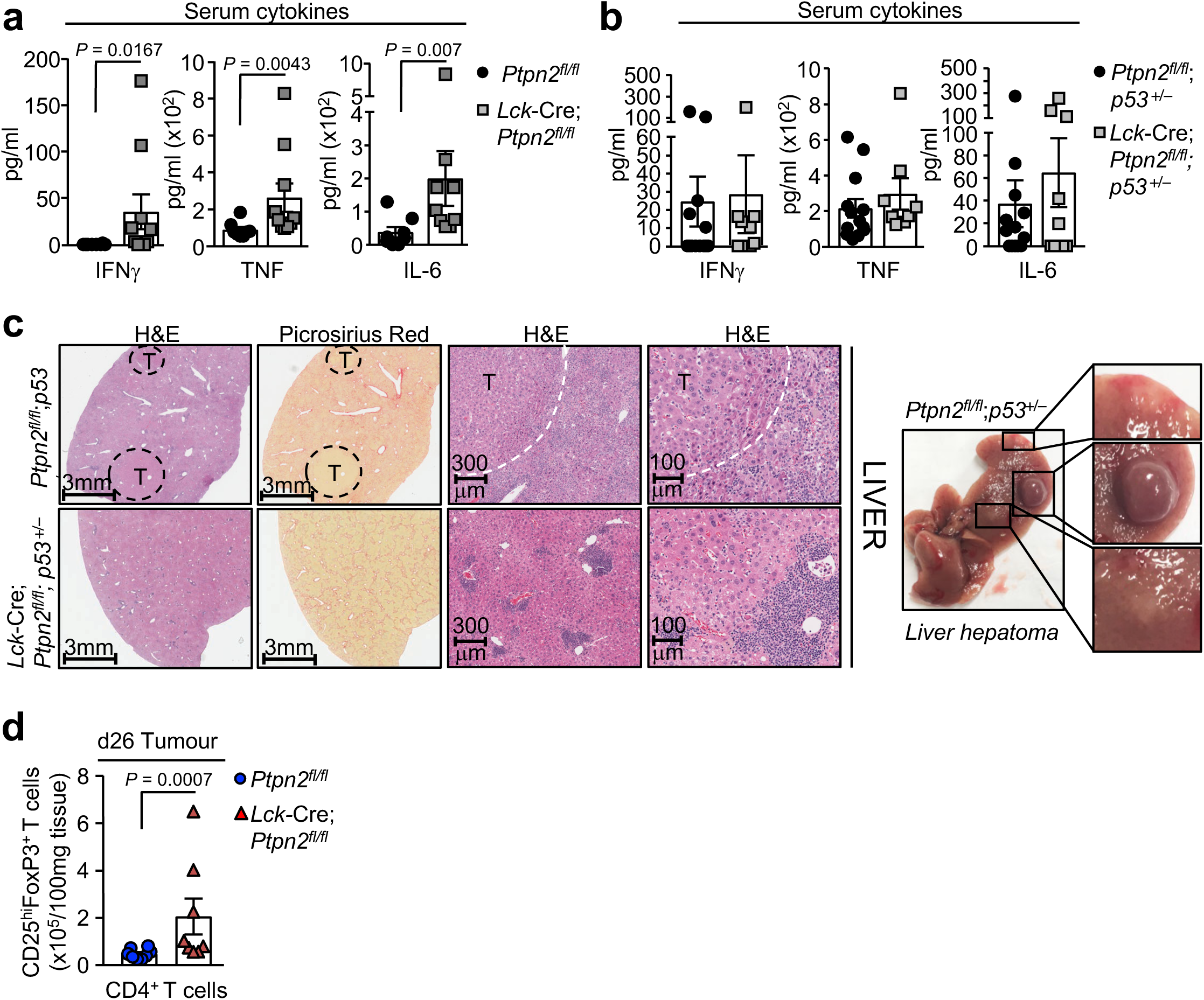
Inflammatory disease in Ptpn2^fl/fl^;p53^+/–^ and *Lck*-Cre;*Ptpn2*^fl/fl^;p53^+/–^ mice. Inflammatory serum cytokines in **a**) *Ptpn2^fl/fl^* versus *Lck*-Cre;*Ptpn2^fl/fl^* mice and **b**) *Ptpn2^fl/fl^*;*p53^+/–^* and *Lck*-Cre;*Ptpn2^fl/fl^;p53^+/–^* mice were determined with the LEGENDplex T_h_ Cytokine Panel™ kit. **b**) Livers from 12 month old *Ptpn2^fl/fl^*;*p53^+/–^* and *Lck*-Cre;*Ptpn2^fl/fl^;p53^+/–^* mice were fixed in formalin and processed for histology. Gross images are shown of *Ptpn2^fl/fl^;p53^+/–^* livers bearing a hepatomas (T). **c**) Day 26 (d26) tumour infiltrating lymphocytes (TILs) from *Ptpn2^fl/fl^* and *Lck*-Cre;*Ptpn2^fl/fl^* mice were stained for CD4, CD25 and intracellular FoxP3 and the proportion of CD4^+^CD25^+^FoxP3^+^ T_regs_ cells determined by flow cytometry. Significance in (a, d) was determined using 2-tailed Mann-Whitney U Test.

**Supplementary figure 3.**
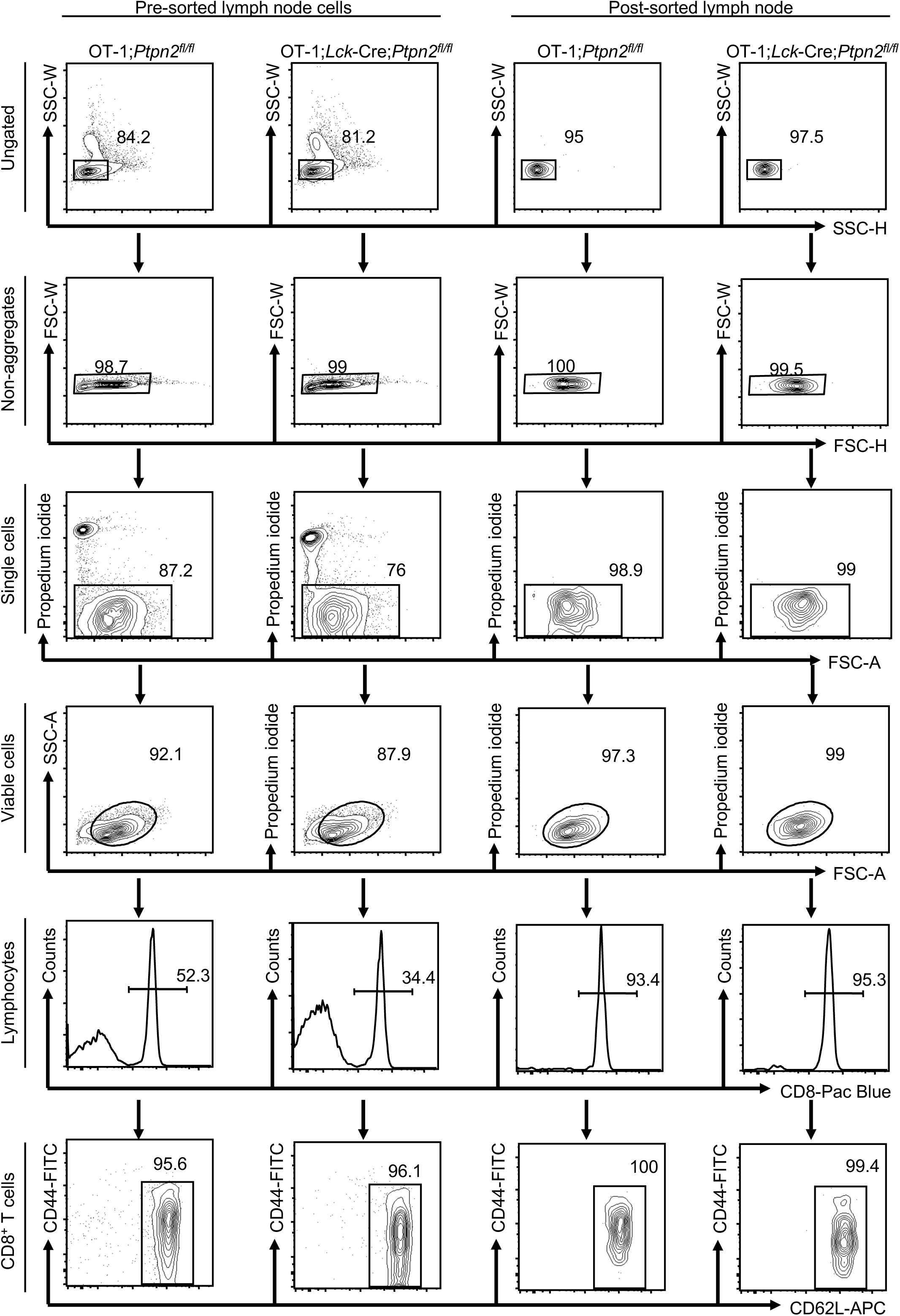
Exemplary FACS-gating strategy. Lymph node cells from OT-1;*Ptpn2^fl/fl^* and OT-1;*Lck*-Cre;*Ptpn2^fl/fl^* mice were stained for CD8, CD44 and CD62L and sorted for CD8^+^CD44^lo^CD62L^hi^ T cells.

**Supplementary figure 5.**
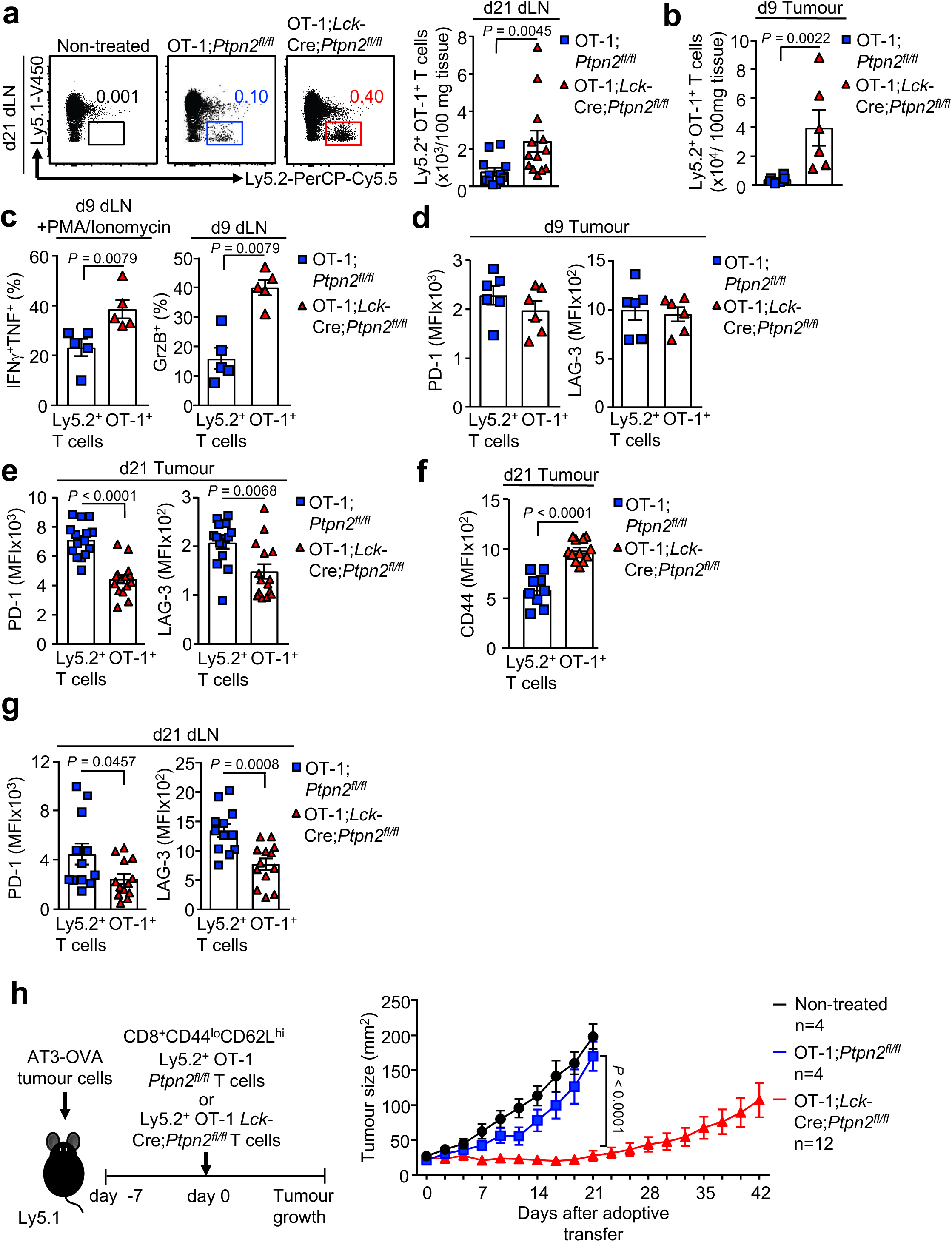
PTPN2 deficiency increases T cell infiltration and activation and decreases T cell exhaustion in a syngeneic model of triple negative breast cancer. AT3-OVA breast cancer cells (1×10^6^) were injected into the fourth inguinal fat pads of female Ly5.1^+^ mice. Seven days after tumour injection FACS-purified naïve CD8^+^CD44^lo^CD62L^hi^ lymph node T cells from Ly5.2^+^ OT-1;*Ptpn2^fl/fl^* (2×10^6^) and Ly5.2^+^ OT-1;*Lck*-Cre;*Ptpn2^fl/fl^* (2×10^6^) mice were adoptively transferred into tumour-bearing Ly5.1 mice. (**a**) 21 days after adoptive transfer lymphocytes were isolated from the draining lymph nodes (dLN) and stained for Ly5.1 and Ly5.2 and Ly5.2^+^ OT-1;*Ptpn2^fl/fl^* and Ly5.2^+^ OT-1;*Lck*-Cre;*Ptpn2^fl/fl^* donor T cell numbers were determined. **b-c**) 9 days after adoptive transfer lymphocytes from mammary tumours or dLN were stained for Ly5.1, Ly5.2 and intracellular IFNγ, TNF and Granzyme B (GrzB) and Ly5.2^+^ OT-1;*Ptpn2^fl/fl^* and Ly5.2^+^ OT-1;*Lck*-Cre;*Ptpn2^fl/fl^* T cell numbers or the proportion of Ly5.2^+^IFNγ^+^TNF^+^ and Ly5.2^+^GrzB^+^ T cells determined. **d-g**) 9 days or 21 days post adoptive transfer lymphocytes from mammary tumours or dLN were stained for Ly5.1, Ly5.2, PD-1 and LAG-3 or CD44 and **d, e, g**) LAG-3 and PD-1 MFIs or **f**) CD44 MFIs in Ly5.2^+^ OT-1;*Ptpn2^fl/fl^* and Ly5.2^+^ OT-1;*Lck*-Cre;*Ptpn2^fl/fl^* T cells determined. **h**) AT3-OVA tumour growth and survival was monitored in female Ly5.1^+^ mice up to 42 days post adoptive transfer. Representative flow cytometry profiles and results (means ± SEM) from at least two independent experiments are shown. Significance in (a-g) was determined using 2-tailed Mann-Whitney U Test. In (h) significance was determined using 2-way ANOVA Test.

**Supplementary figure 5.**
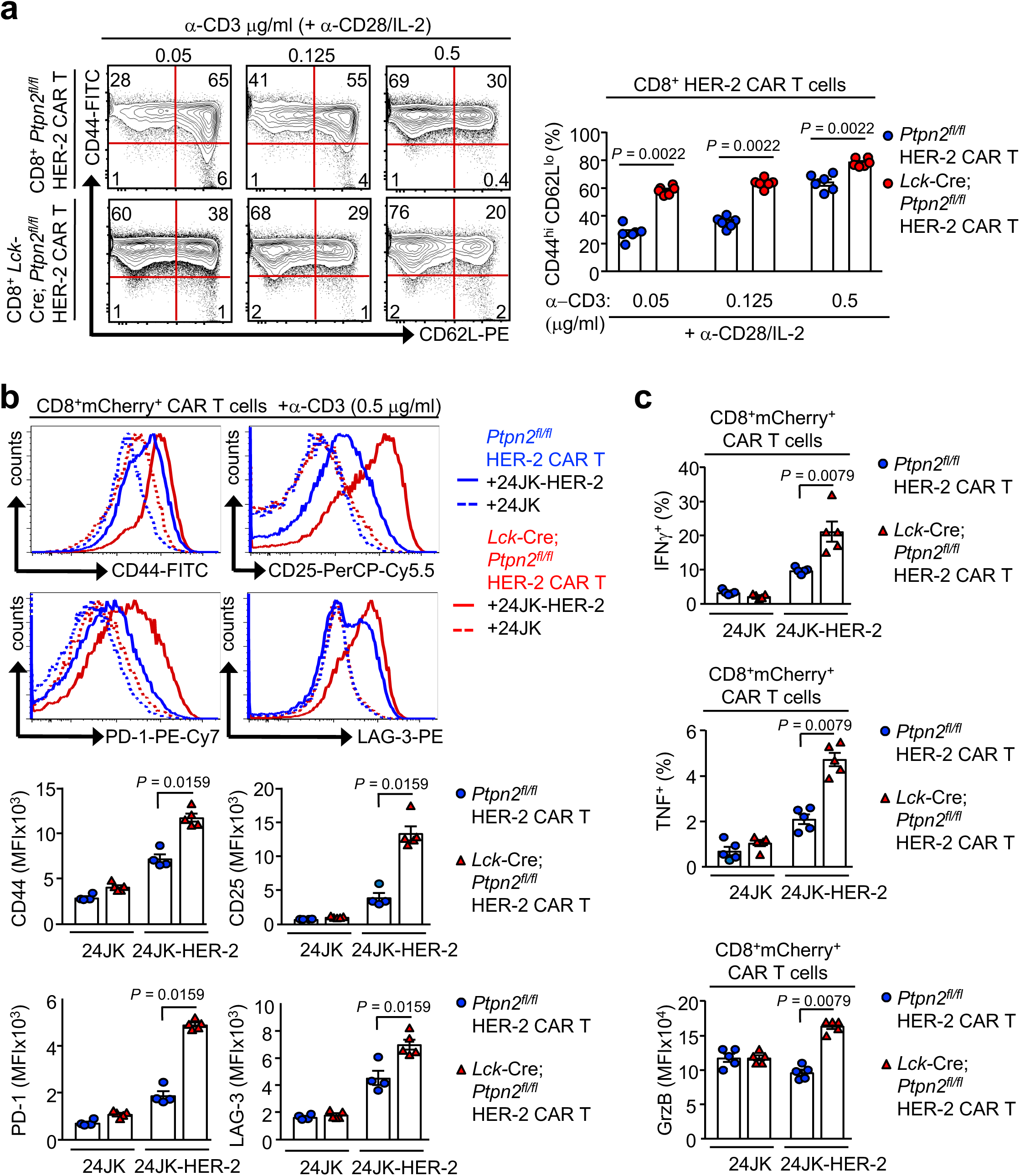
PTPN2 deficiency enhances the generation and the antigen-induced activation of CAR T cells. **a**) *Ptpn2^fl/fl^* versus *Lck*-Cre;*Ptpn2^fl/fl^* HER-2 CAR T cells were generated with varying concentrations of α-CD3 (0.05, 0,125 and 0.5 μg/ml) in the presence of α-CD28 (0.5 μg/ml) and IL-2 (2 ng/ml). After 6 days in culture HER-2 CAR T cells were stained for CD8, CD44 and CD62L and the generation of effector/memory (CD44^hi^CD62L^lo^) CAR T cells determined by flow cytometry. **b-c**) *Ptpn2^fl/fl^* versus *Lck*-Cre;*Ptpn2^fl/fl^* HER-2 CAR T cells were incubated with HER-2 expressing 24JK sarcoma cells (24JK-HER-2) and HER-2 negative 24JK sarcoma cells. Cells were stained for **b**) CD8, CD25, CD44, PD-1 and LAG-3 and **c**) CD8, intracellular IFNγ, TNF and Granzyme B (GrzB) and **b**) CD25, CD44, PD-1, LAG-3 MFIs and **c**) the proportion of IFNγ^+^ and TNF^+^ CAR T cells and GrzB MFIs were determined by flow cytometry. Significance in (a-c) was determined using 2-tailed Mann-Whitney U Test.

**Supplementary figure 6.**
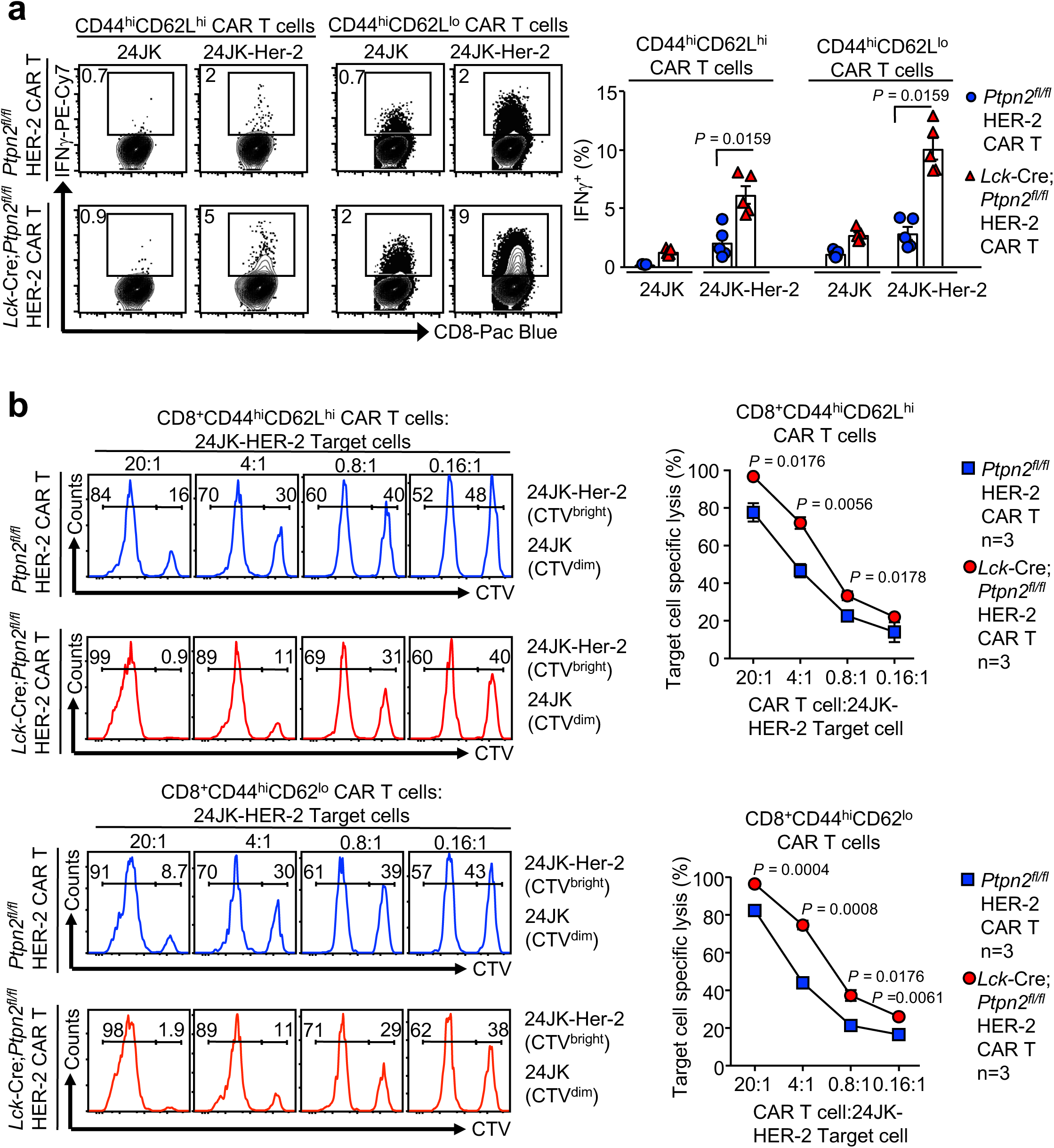
PTPN2 deficiency enhances the antigen-induced cytotoxic activity of CAR T cells. **a**) *Ptpn2^fl/fl^* versus *Lck*-Cre;*Ptpn2^fl/fl^* HER-2 CAR T cells were incubated with 24JK-HER-2 or 24JK sarcoma cells. Cells were stained for CD8, CD44, CD62L and intracellular IFNγ, and the proportion of IFNγ^+^ CAR T cells determined by flow cytometry. **b**) FACS-purified CD8^+^ central memory (CD44^hi^CD62L^hi^) and effector/memory (CD44^hi^CD62L^lo^) *Ptpn2^fl/fl^* versus *Lck*-Cre;*Ptpn2^fl/fl^* HER-2 CAR T cells were incubated with 5 mM CTV-labelled (CTV^bright^) 24JK-HER-2 and 0.5 mM CTV-labelled (CTV^dim^) 24JK sarcoma cells. Antigen-specific target cell lysis (24JK-HER-2 versus 24JK response) was monitored for the depletion of CTV^bright^ 24JK-HER-2 cells by flow cytometry. Representative flow cytometry profiles and results (means ± SEM) from at three independent experiments are shown. Significance in (a) was determined using 2-tailed Mann-Whitney U Test. Significance in (b) was determined using 2-tailed Student’s t test.

**Supplementary figure 7.**
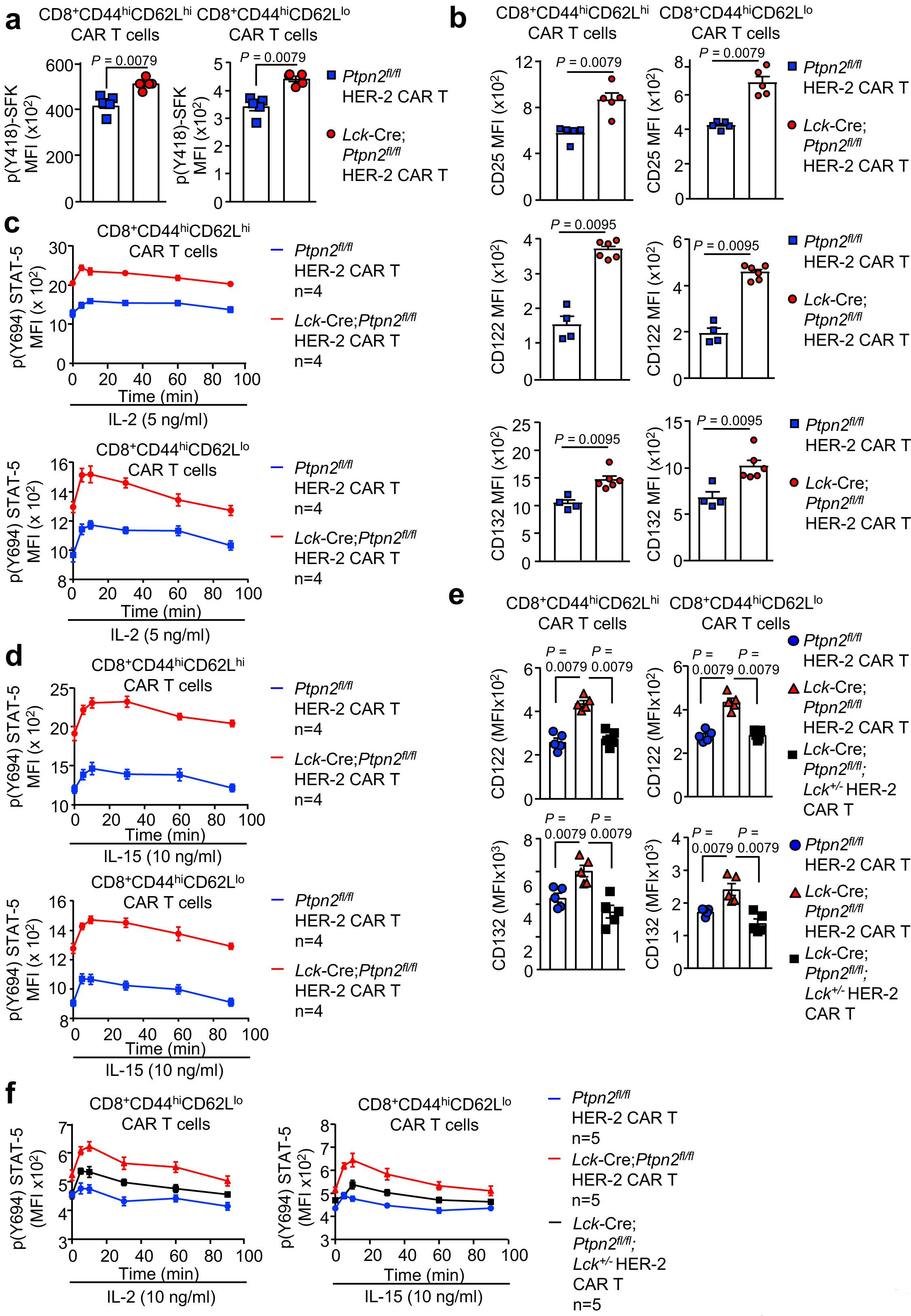
PTPN2 deficiency increases SFK and IL-2/15-induced STAT5 signaling in CD8^+^ HER-2 CAR T cells. **a**) *Ptpn2^fl/fl^* versus *Lck*-Cre;*Ptpn2^fl/fl^* HER-2 CAR T cells were stained for CD8 and intracellular p(Y418)-SFK and p(Y418)-SFK mean fluorescence intensity (MFI) was determined by flow cytometry. **b**) HER-2-specific *Ptpn2^fl/fl^* versus *Lck*-Cre;*Ptpn2^fl/fl^* CAR T cells were incubated with plate-bound α-CD3 and stained for CD8, CD62L, CD44, CD25, CD122 and CD132 and CD25, CD122 and CD132 MFIs on CD8^+^CD44^hi^CD62L^hi^ CAR T cells versus CD8^+^CD44^hi^CD62L^lo^ CAR T cells were determined by flow cytometry. **c-d**) *Ptpn2^fl/fl^* versus *Lck*-Cre;*Ptpn2^fl/fl^* HER-2 CAR T cells were incubated with plate-bound α-CD3 and stimulated with murine recombinant **c**) IL-2 and **d**) IL-15 for the indicated time points. Cells were stained for CD8, CD62L, CD44 and intracellular p(Y694)-STAT5 MFIs on versus CD8^+^CD44^hi^CD62L^hi^ CAR T cells versus CD8^+^CD44^hi^CD62L^lo^ were determined by flow cytometry. **e**) *Ptpn2^fl/fl^*, *Lck*-Cre;*Ptpn2^fl/fl^* and *Lck*-Cre;*Ptpn2^fl/fl^*;*Lck^+/-^* HER-2 CAR T cells were incubated with plate-bound α-CD3 and stained for CD8, CD44, CD62L, CD122 and CD132 and CD122 and CD132 MFIs on CD8^+^CD44^hi^CD62L^lo^ versus CD8^+^CD44^hi^CD62L^hi^CAR T cells were determined by flow cytometry. **f**) *Ptpn2^fl/fl^* versus *Lck*-Cre;*Ptpn2^fl/fl^* or *Lck*-Cre;*Ptpn2^fl/fl^*;*Lck^+/-^* HER-2 CAR T cells were incubated with plate-bound α-CD3 and stimulated with murine recombinant IL-2 and IL-15 for the indicated time points. Cells were stained for CD8, CD62L, CD44 and intracellular p(Y694)-STAT5 MFIs on CD8^+^CD44^hi^CD62L^lo^ CAR T cells were determined by flow cytometry. Representative results (means ± SEM) are shown from two independent experiments. In (a-b, e) significance was determined using 2-tailed Mann-Whitney U Test.

**Supplementary figure 8.**
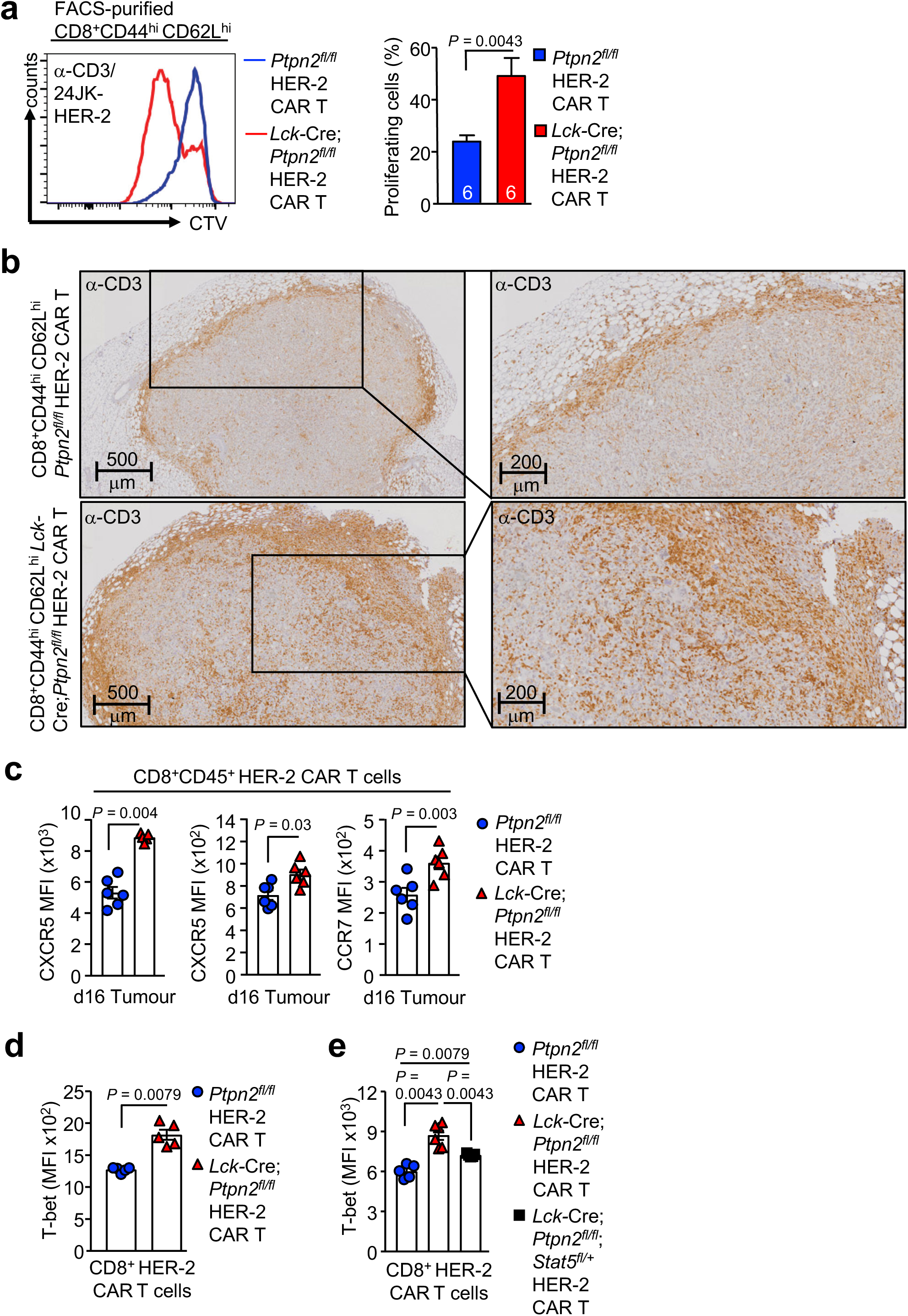
PTPN2 deficiency enhances CAR T cell proliferation and tumour infiltration. **a**) CD8^+^CD44^hi^CD62L^hi^ HER-2 CAR T cells generated from *Ptpn2^fl/fl^* versus *Lck*-Cre;*Ptpn2^fl/fl^* splenocytes were stimulated with plate-bound α-CD3 and subsequently labelled with CTV and incubated with 24JK-HER-2 cells and proliferation was determined by flow cytometry. **b**) HER-2-E0771 tumours isolated from HER-2 TG mice on day 10 after adoptive *Ptpn2^fl/fl^* versus *Lck*-Cre;*Ptpn2^fl/fl^* HER-2 CAR T cell transfer were analysed for CD3^+^ T cell infiltrates by immunohistochemistry. **c**) HER-2-E0771 cells (2×10^5^) were injected into the fourth inguinal mammary fat pads of female HER-2 transgenic (TG) mice. Seven days after tumour injection HER-2 TG mice received total body irradiation (4 Gy) followed by the adoptive transfer of 6×10^6^ FACS-purified CD8^+^CD62L^hi^CD44^hi^ central memory HER-2 CAR T cells generated from *Ptpn2^fl/fl^* or *Lck*-Cre;*Ptpn2^fl/fl^* splenocytes. Mice were injected with IL-2 (50,000 IU/day) on days 0-4 after adoptive CAR T cell transfer. Lymphocytes were isolated from the tumours at day 16 post adoptive transfer and stained for CD45, CD8, CXCR3, CXCR5 and CCR7 and CXCR3, CXCR5 and CCR7 MFIs were determined by flow cytometry. **d-e**) CD8^+^ HER-2 CAR T cells generated from *Ptpn2^fl/fl^*, *Lck*-Cre;*Ptpn2^fl/fl^* or *Lck*-Cre;*Ptpn2^fl/^*^fl^;*Stat5^fl/+^* splenocytes were stained for intracellular T-bet and T-bet MFIs determined by flow cytometry. Representative flow cytometry profiles and results (means ± SEM) are shown from two independent experiments. In (a, c, d, e) significance was determined using 2-tailed Mann-Whitney U Test.

**Supplementary figure 9.**
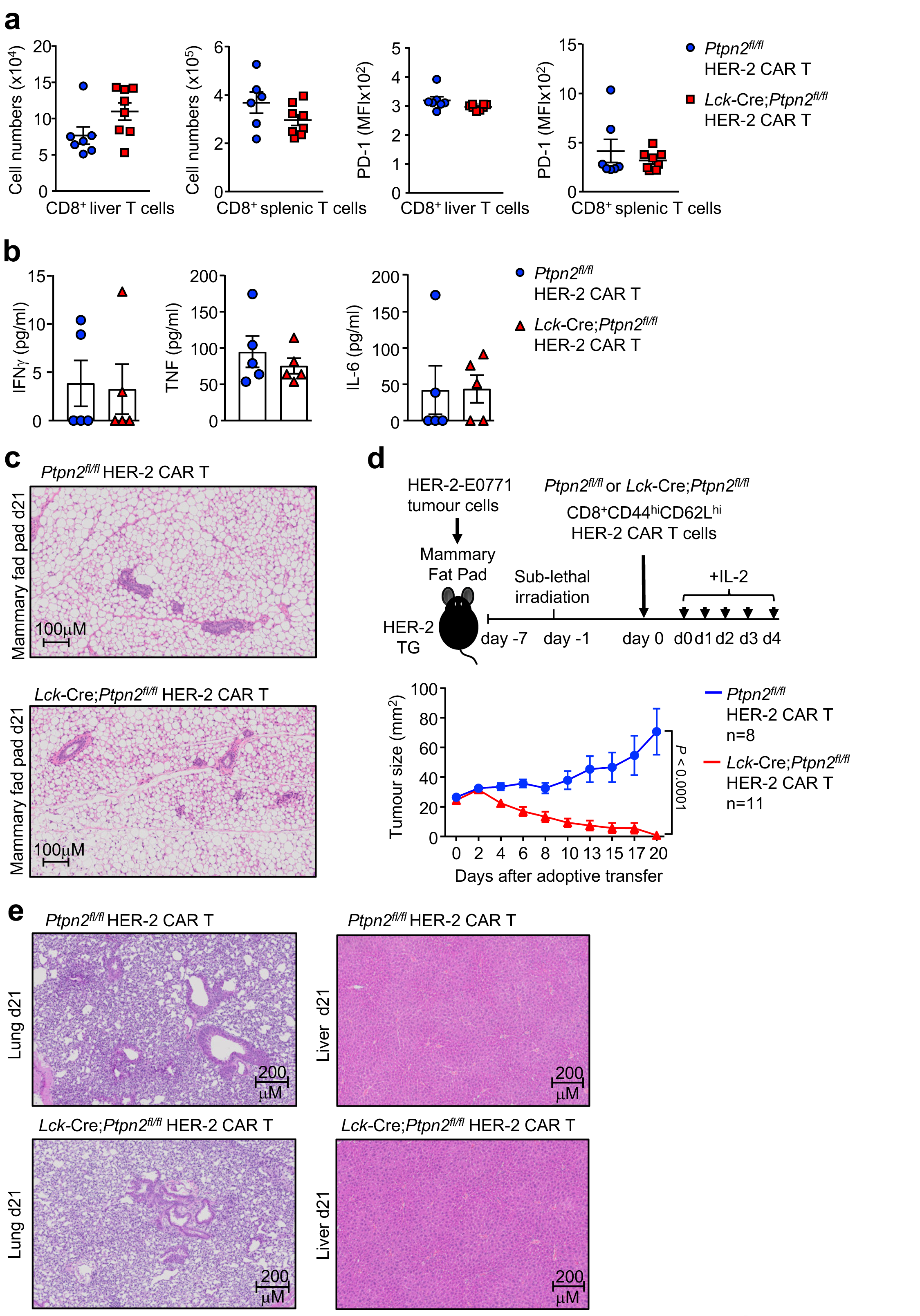
PTPN2 deficiency in CAR T cells does not result in systemic inflammation and collateral tissue damage. HER-2-E0771 cells (2×10^5^) were injected into the fourth inguinal mammary fat pads of female HER-2 TG mice. Seven days after tumour injection HER-2 TG mice received total body irradiation (4 Gy) followed by the adoptive transfer of 6×10^6^ FACS-purified CD8^+^CD44^hi^CD62L^hi^ central memory HER-2 CAR T cells generated from *Ptpn2^fl/fl^* or *Lck*-Cre;*Ptpn2^fl/fl^* splenocytes. Mice were injected with IL-2 (50,000 IU/day) on days 0-4 after adoptive CAR T cell transfer. **a**) Lymphocytes isolated from the spleens and livers of HER-2 TG recipient mice were stained for CD3 and CD8 and CD3^+^CD8^+^ donor CAR T cell numbers were determined by flow cytometry 21 days after adoptive CAR T cell transfer. **b**) Inflammatory serum cytokines in HER-2 TG recipient mice were determined with the LEGENDplex T_h_ Cytokine Panel™ kit. **c**) The tumour-free contralateral fourth inguinal mammary fat pads were fixed in formalin at 21 days post CAR T cell transfer and processed for histological assessment (hematoxylin and eosin: H&E) monitoring for tissue architecture and lymphocytic infiltrates. **d**) HER-2-E0771 breast cancer cells (2×10^5^) were injected into the fourth inguinal mammary fat pads of female HER-2 TG mice. Seven days after tumour injection HER-2 TG mice received total body irradiation (4 Gy) followed by the adoptive transfer of 20×10^6^ FACS-purified CD8^+^CD44^hi^CD62L^hi^ central memory HER-2 CAR T cells generated from *Ptpn2^fl/fl^* or *Lck*-Cre;*Ptpn2^fl/fl^* splenocytes. Mice were injected with IL-2 (50,000 IU/day) on days 0-4 after adoptive CAR T cell transfer and tumour growth was monitored. **e**) Lungs and livers were fixed in formalin at 21 days post CAR T cell transfer and processed for histological assessment (hematoxylin and eosin: H&E) monitoring for tissue architecture and lymphocytic infiltrates. Representative results (means ± SEM) from at least two independent experiments are shown. In (d) significance was determined using 2-way ANOVA Test.

**Supplementary figure 10.**
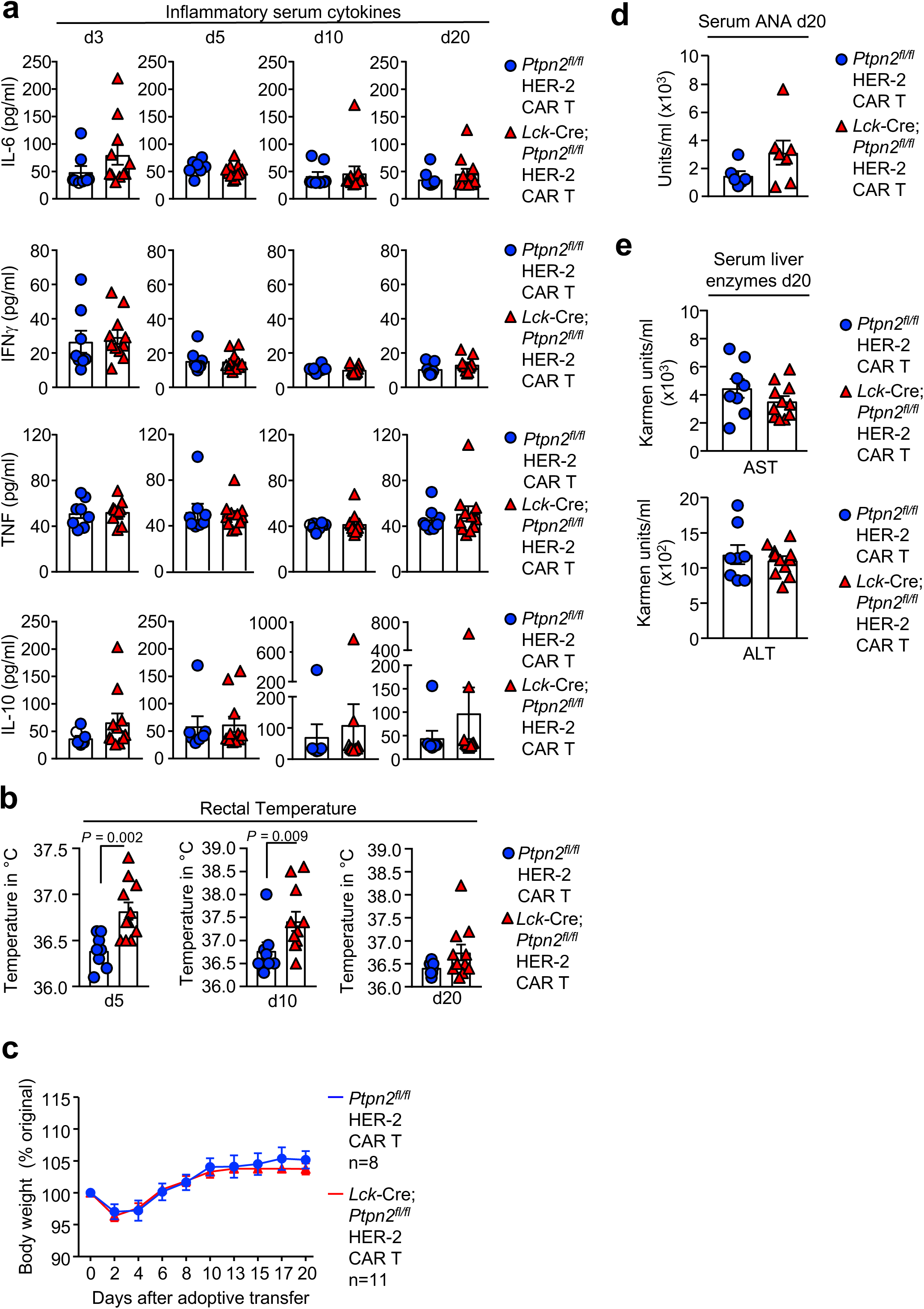
PTPN2 deficiency in CAR T cells does not result in autoimmunity. HER-2-E0771 breast cancer cells (2×10^5^) were injected into the fourth inguinal mammary fat pads of female HER-2 TG mice. Seven days after tumour injection HER-2 TG mice received total body irradiation (4 Gy) followed by the adoptive transfer of 20×10^6^ FACS-purified CD8^+^CD44^hi^CD62L^hi^ central memory HER-2 CAR T cells generated from *Ptpn2^fl/fl^* or *Lck*-Cre;*Ptpn2^fl/fl^* splenocytes. Mice were injected with IL-2 (50,000 IU/day) on days 0-4 after adoptive CAR T cell transfer. **a**) Serum cytokines were determined with the BD CBA Mouse Inflammation Kit™. **b**) Body core temperatures were measured using a mouse rectal probe and **c**) body weights monitored. **d**) Serum anti-nuclear antibodies (ANA) were measured using a mouse anti-nuclear antibodies Ig’s (total IgA+G+M) ELISA Kit. **e**) Serum ALT and AST activities were determined using a Transaminase II Kit.

**Supplementary figure 11.**
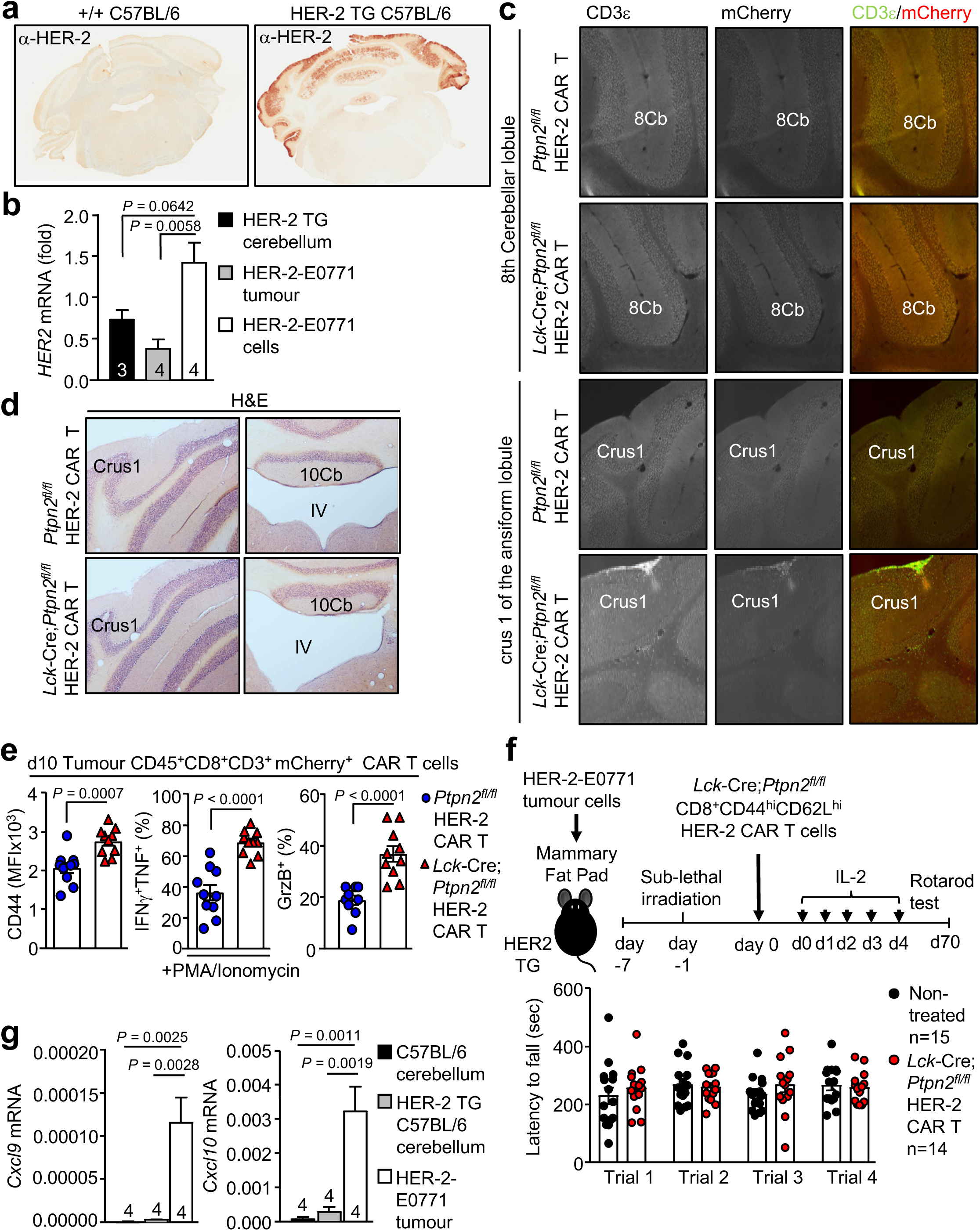
PTPN2 deficiency does not result in marked CAR T cell cerebellar infiltration and tissue damage. **a**) Cerebella from +/+ C57BL/6 and HER-2 TG C57BL/6 mice were processed for α-HER-2 immunohistochemistry. **b**) *HER-2* gene expression in cerebella from HER-2 TG mice, E0771-HER-2 tumours or HER-2-E0771 cells were assessed by quantitative real time PCR. **c**) Cerebella from HER-2 TG C57BL/6 mice administered *Ptpn2^fl/fl^* or *Lck*-Cre;*Ptpn2^fl/fl^* mCherry+ HER-2 CAR T cells were processed for α-CD3ε immunohistochemistry 10 days after the transfer of HER-2 CAR T cells. CD3ε^+^mCherry^+^ CAR T cells were not detected in the cerebellar lobules, including the 8^th^ cerebellar lobule (8Cb). Some CD3ε^+^mCherry^+^ staining was evident adjacent to the crus 1 of the ansiform lobule (Crus1) in mice treated with PTPN2-deficienct HER-2 CAR T cells. Representative images for 3 mice per genotype are shown. **d**) Cerebella from HER-2 TG mice administered *Ptpn2^fl/fl^* or *Lck*-Cre;*Ptpn2^fl/fl^* HER-2 CAR T cells were processed for histology (hematoxylin and eosin: H&E) 10 days after the adoptive transfer of HER-2 CAR T cells to assess gross tissue architecture. **e**) *Ptpn2^fl/fl^* versus *Lck*-Cre;*Ptpn2^fl/fl^* HER-2 CAR T cells isolated from HER-2-E0771 tumours 10 days post adoptive transfer were stained for CD45, CD8, CD3, CD44 and intracellular IFNγ, TNF and GrzB and CD44 MFIs and the proportion of IFNγ^+^TNF^+^ and GrzB^+^ CAR T cells determined by flow cytometry. **f**) HER-2 TG mice administered *Lck*-Cre;*Ptpn2^fl/fl^* HER-2 CAR T cells were subjected to a rotarod test 50 days post tumour clearance and the latency to fall determined in consecutive trials. **g**) *Cxcl9 and Cxcl10* gene expression in cerebella from C57Bl/6 and HER-2 TG mice and E0771-HER-2 tumours were assessed by quantitative real time PCR. Representative results (means ± SEM) from at least two independent experiments are shown. In (c, g) significance was determined using 1-way ANOVA Test. In (e) significance was determined using 2-tailed Mann-Whitney U Test.

**Supplementary figure 12.**
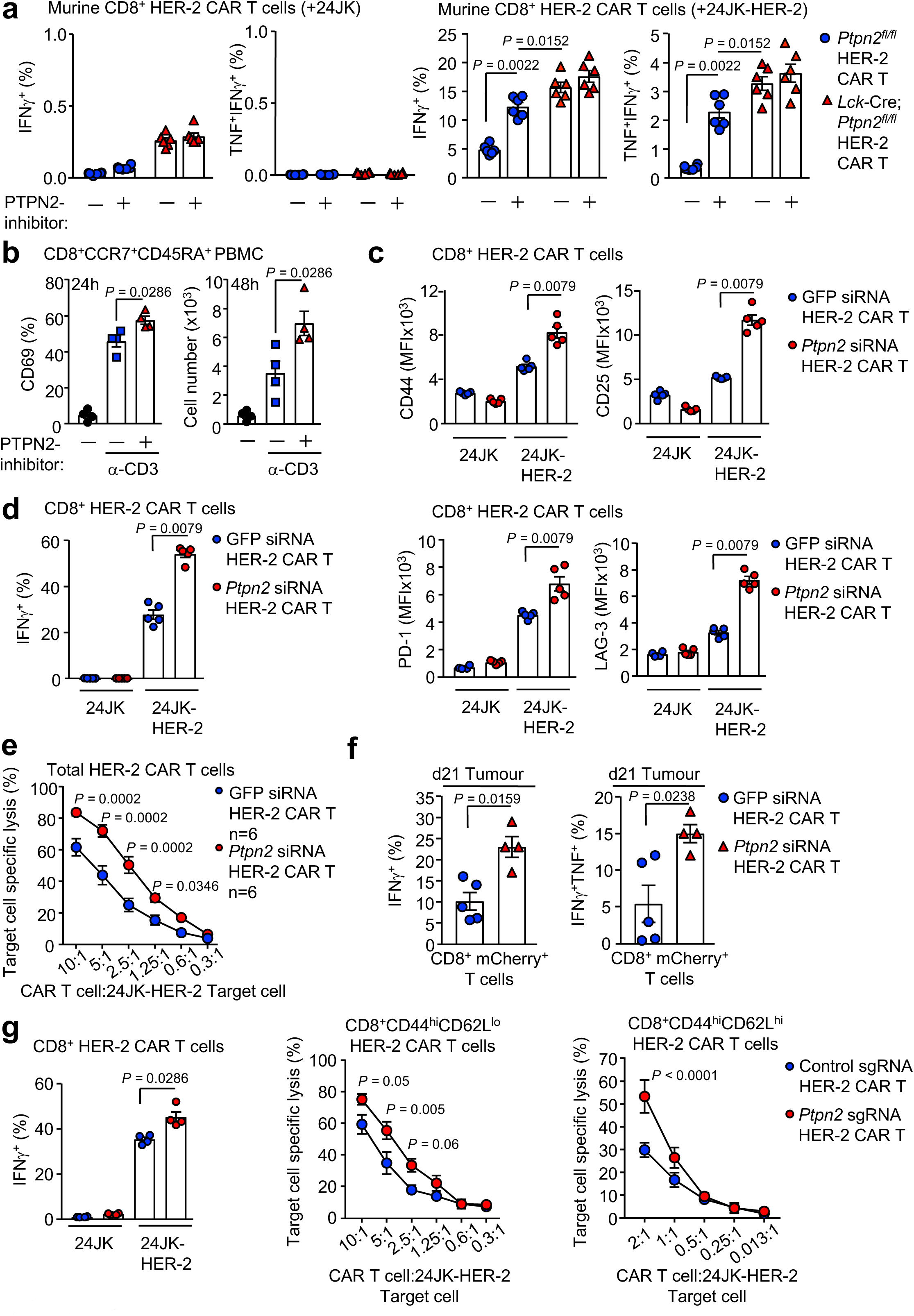
Targeting Ptpn2 with siSTABLE™ siRNAs or using CRISPR-Cas9 RNP enhances tumour-specific CAR T cell responses. **a**) *Ptpn2^fl/fl^* versus *Lck*-Cre;*Ptpn2^fl/fl^* CD8^+^ HER-2 CAR T cells were treated with PTPN2-inhibitor (+) or vehicle (-) followed by incubation with 24JK-HER-2 versus 24JK sarcoma cells. Cells were stained for CD8 and intracellular IFNγ and TNF and the proportion of CD8^+^IFNγ^+^ and CD8^+^IFNγ^+^TNF^+^ CAR T cells determined by flow cytometry. **b**) Human PBMCs were pretreated with PTPN2-inhibitor (+) or vehicle (-) and stimulated with α-CD3. Cells were stained for CD8, CCR7, CD45RA and CD69 and MFIs for CD69 on CD8^+^CCR7^+^CD45RA^+^ T cells and CD8^+^CCR7^+^CD45RA^+^ T cell numbers were determined by flow cytometry. **c**) HER-2 CAR T cells transfected with GFP versus *Ptpn2* siSTABLE™ siRNAs were incubated with 24JK-HER-2 versus 24JK sarcoma cells. Cells were stained for CD8, CD25, CD44, PD-1 and LAG-3 and CD25, CD44, PD-1 and LAG-3 MFIs were determined by flow cytometry. **d**) HER-2 CAR T cells transfected with GFP versus *Ptpn2* siSTABLE™ siRNAs were incubated with 24JK-HER-2 versus 24JK sarcoma cells. Cells were stained with fluorochrome-conjugated antibodies for CD8 and intracellular IFNγ, and the proportion of IFNγ^+^ CAR T cells determined by flow cytometry. **e**) HER-2 CAR T cells transfected with GFP versus *Ptpn2* siSTABLE™ siRNAs were incubated with 5 mM CTV-labelled (CTV^bright^) 24JK-HER-2 and 0.5 mM CTV-labelled (CTV^dim^) 24JK sarcoma cells. Antigen-specific target cell lysis (24JK-HER-2 versus 24JK response) was monitored for the depletion of CTV^bright^ 24JK-HER-2 cells by flow cytometry. **f**) CD8^+^ mCherry^+^ CAR T cells isolated from HER-2-E0771 tumours 21 days post adoptive transfer were stained for CD8 and intracellular IFNγ and TNF and the proportion of IFNγ^+^ and IFNγ^+^TNF^+^ CAR T cells determined by flow cytometry. **g-h**) HER-2 CAR T cells were transfected with control versus *Ptpn2* sgRNAs plus Cas9 using the Lonza 4D-Nucleofector to delete PTPN2 by CRISPR-Cas9 RNP. **g**) Control and PTPN2 deleted HER-2 CAR T cells were incubated with 24JK-HER-2 versus 24JK sarcoma cells and stained for CD8 and intracellular IFNγ and the proportion of IFNγ^+^ CAR T cells determined by flow cytometry. **h**) Alternatively, control and PTPN2-deleted HER-2 CAR T cells were incubated with 5 mM CTV-labelled (CTV^bright^) 24JK-HER-2 cells and 0.5 mM CTV-labelled (CTV^dim^) 24JK sarcoma cells. Antigen-specific target cell lysis (24JK-HER-2 versus 24JK response) was assessed by monitoring for the depletion of CTV^bright^ 24JK-HER-2 cells by flow cytometry. Representative results (means ± SEM) from at least two independent experiments are shown. In (a-d, f) significance was determined using 2-tailed Mann-Whitney U Test. In (e, g) significance was determined using 2-way ANOVA Test.

**Table S1.** Pathology in *Ptpn2^fl/fl^;p53* ^+/-^ mice

| Mouse ID | Hepatoma | Sarcoma/<br>Carcinoma | Thymoma | Lymphoma | Splenomegaly | B1 cell<br>leukemia | Thymic DP T<br>cell leukemia | Splenic/Hepatic<br>DP T cell leukemia |
| --- | --- | --- | --- | --- | --- | --- | --- | --- |
| #1 | No | No | Yes | No | No | No | Yes | Yes |
| #2 | No | No | No | Yes | Yes | Yes | No | No |
| #4 | No | No | No | No | No | Yes | No | No |
| #6 | No | No | No | No | Yes | Yes | No | No |
| #7 | No | Yes | No | No | No | No | Yes | Yes |
| #8 | No | No | No | No | No | No | Yes | No |
| #14 | Yes | No | No | No | Yes | No | No | No |
| #15 | No | No | No | No | Yes | Yes | No | No |
| #18 | No | Yes | No | No | No | No | No | No |
| #20 | No | No | Yes | No | No | No | Yes | Yes |
| #23 | No | No | Yes | No | No | No | Yes | Yes |
| #27 | No | No | No | No | Yes | Yes | No | No |
| #30 | No | Yes | No | No | No | No | No | No |
| #32 | No | No | No | No | Yes | Yes | No | No |
| #34 | Yes | No | No | No | No | No | No | No |

